# Mapping brain networks underlying abstract perceptual decision formation and its EEG signatures

**DOI:** 10.64898/2026.10.29.762507

**Authors:** Abdoreza Asadpour, Natalie A. Steinemann, Katerina Kalou, Simon P. Kelly, Redmond G. O’Connell, KongFatt Wong-Lin

## Abstract

Perceptual decisions have been proposed to involve a common mechanism of evidence accumulation operating across sensory modalities and motor contexts. In human EEG, the centroparietal positivity (CPP) tracks evidence accumulation in such a supramodal motor-independent manner, but its cortical sources are uncertain. Here we combined fMRI and EEG, acquired in separate sessions of the same supramodal change-detection paradigm, with individualized finite element head modeling to identify decision-related networks. A conjunction analysis across visual and auditory modalities and across motor and non-motor required tasks revealed consistent activation of bilateral postcentral, supramarginal, and superior parietal lobules, precentral gyrus, central operculum, middle frontal gyrus, and cerebellum. Forward modeling of fMRI-defined sources indicated that parietal–opercular regions were the dominant contributors to the response-locked and non-motor-required peak-locked CPP signal. These results implicate a network of brain areas in supramodal, motor-independent target-detection decisions, whose specific functional roles can be interrogated in more complex decision tasks in future.

## 1 Introduction

Perceptual decision-making is understood to involve the processes of sensory evidence encoding, decision formation via temporal accumulation of evidence, and motor action when required (Shadlen & Kiani, 2013). Prior studies across various species have identified distributed neural correlates of the aforementioned processes during perceptual decision-making (Brown et al., 2026; Gherman et al., 2024; Hanks & Summerfield, 2017; Khilkevich et al., 2024; Meshulam et al., 2025; O’Connell et al., 2018; Okazawa & Kiani, 2023).

In particular, the evidence accumulation process has been a key guiding light when searching for the neural basis of decision formation (Gold & Shadlen, 2007; Hanks & Summerfield, 2017; Heekeren et al., 2004; O’Connell et al., 2018). While evidence accumulation dynamics have long been observed in neural circuits associated with the planning of choice-reporting motor actions (Gold & Shadlen, 2007), many studies in recent years have sought to identify motor-independent processes that could play a more central, flexible, mediating role between any relevant sensory information and potential actions in the face of changing contexts or demands (Freedman & Assad, 2016; O’Connell et al., 2018). In human intracranial electrophysiology, motor-independent evidence accumulation dynamics have been observed at distributed brain sites (Gherman et al., 2024; Xie et al., 2024). In non-invasive electrophysiology (scalp EEG), a centroparietal positivity (CPP) has been found to exhibit evidence accumulation dynamics regardless of either sensory modality or motor requirements (O’Connell et al., 2012; Twomey et al., 2016), to correlate with choice accuracy (O’Connell & Kelly, 2021), and to temporally precede motor preparation (Kelly & O’Connell, 2013) all consistent with a central, intermediate decision process.

fMRI studies seeking to localise such a central process have found activation in the dorsolateral prefrontal cortex, intraparietal sulcus, inferior frontal cortex, and right insula in common across different sensory modalities and/or effectors (Ho et al., 2012; Liu & Pleskac, 2011; Tosoni et al., 2008). In the absence of sufficient temporal resolution, these imaging studies used assumed amplitude differences across perceptual difficulty levels as criteria for identifying accumulation, but such differences strongly depend on unmeasured post-decision dynamics (Twomey et al., 2016). More recently, Wongtrakun et al. (2025) further examined the CPP’s cortical origins using CPP-informed neuroimaging, and found that activation in the intraparietal sulcus covaried with single-trial CPP buildup rate preceding a motor action. However, it remains unclear what regions activate commonly across motor requirements and sensory modalities in the ways observed for the CPP. If CPP really represents such “abstract” decision information, where is the source of the brain circuit that generates it?

In this study, we sought to localise the source of the CPP using a distinct approach involving subject-specific finite-element volume conduction models with individually digitised electrode positions and fMRI-derived spatial priors, and the same paradigm design as used originally to identify the motor-independent, supramodal CPP (O’Connell et al., 2012). Specifically, sensory modalities and motor requirements were manipulated across blocks in which targets were defined as changes in either visual contrast or audio volume from a continuously-presented baseline, and which required either immediate detection reports or critically, a later count of covertly detected targets. Once we mapped brain regions exhibiting common BOLD-signal activation during target-detection across these task conditions, we further interrogated those regions as candidate generators of the CPP measured in EEG data recorded from the same participants performing the same tasks. We found that, among a wider set of commonly-activated areas, the bilateral parietal cortex and bilateral opercular-frontal regions emerged as the most likely sources to produce the centroparietal EEG component associated with supramodal evidence-accumulation.

## 2 Materials and Methods

Data were collected in two experiments involving a total of 45 human participants, who all gave informed consent for the procedures, which were approved by the Trinity College Dublin School of Psychology Ethics Committee. Experiment I, which employed simultaneous EEG-fMRI data collection, included 14 participants (mean age: 24.3 ± 3.5 years, 6 females). Experiment II, which involved separate EEG and fMRI sessions, included 31 participants (mean age: 25.7 ± 4.1 years, 15 females).

### 2.1 Behavioral task paradigms

Participants performed six different conditions of a continuous detection task in separate experimental blocks. During all conditions, participants continuously monitored an oscillating sensory stimulus for changes in intensity, which occurred at pseudo-randomly distributed inter-target-intervals of 4, 6, or 8 s. Participants were instructed to maintain fixation on a central square (5x5 pixels) and did not receive any feedback after the offset of target events.

Visual stimuli consisted of a flickering annular pattern of alternating dark and light radial segments, with two cycles per quadrant presented on a gray background. The annulus had an inner radius of 1.14° and an outer radius of 2.29°, and oscillated at a frequency of 21.25 Hz. During the ITI, the stimulus had a baseline contrast of 70%. During “visual down” blocks, targets were defined by linear contrast decreases from 70% to 40% over 1.6s and a return to 70% in the subsequent 0.8 s (Fig 1A). During “visual-up” blocks, an identical annulus instead linearly rose to 100% contrast during targets.

**Figure 1.**
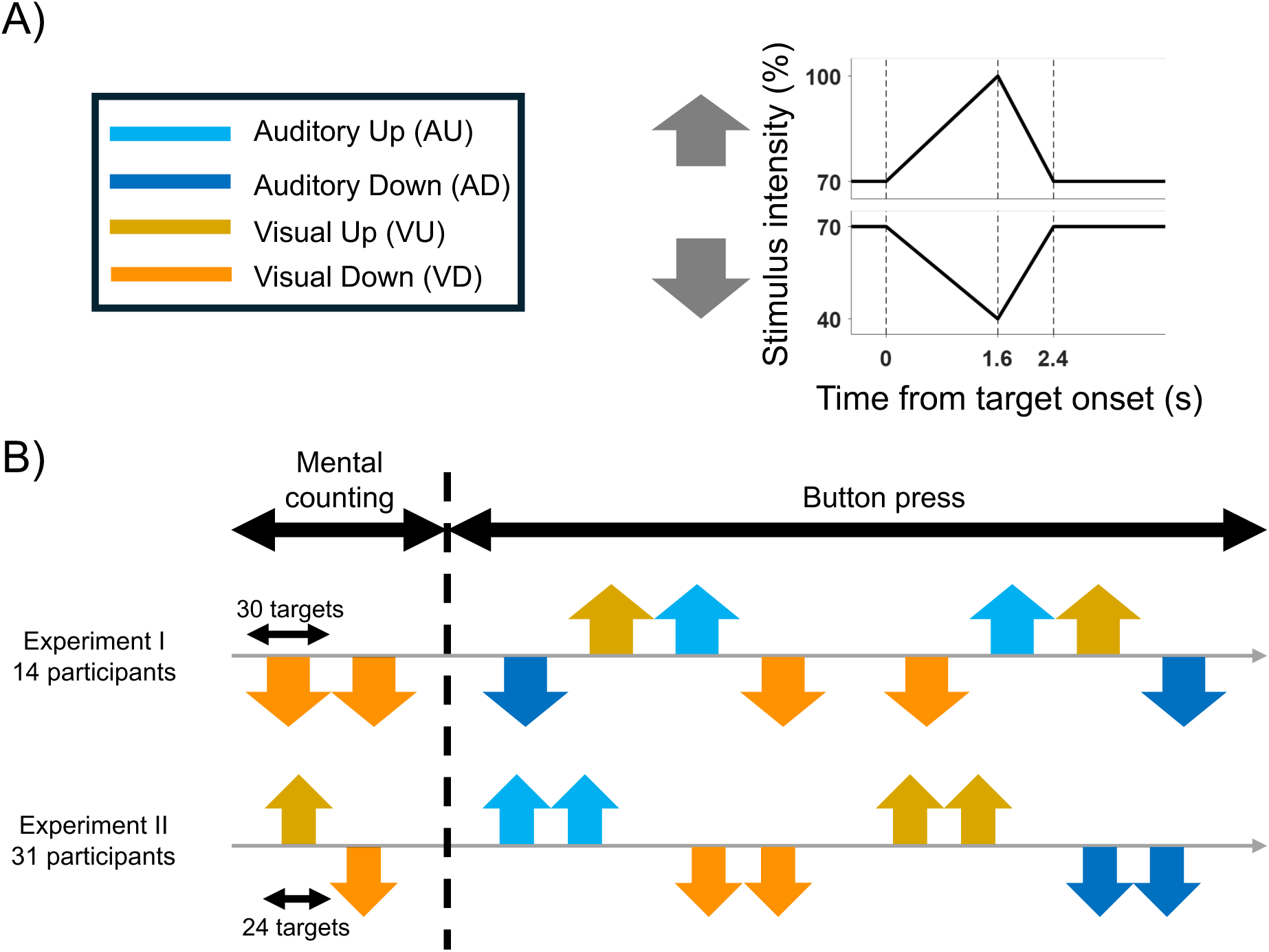
Experimental design for identifying supramodal decision-making brain regions. A) Participants monitored continuous auditory or visual stimuli for ±30% intensity changes (auditory up/down, visual up/down). B) Data were recorded in two separate experiments. In both Experiment I (simultaneous EEG/fMRI) and Experiment II (EEG and fMRI in separate sessions), participants first performed two blocks of a mental counting task, in with they reported the total number of targets detected in an experimental block after the fact, followed by eight blocks of an immediate response task, in which the detection of a target was reported immediately using a button press.

During two auditory conditions, participants monitored a continuous 500Hz tone envelope-modulated at 30Hz, for targets that were defined by linear increases or decreases in auditory volume. The auditory baseline volume was held around 70 dB, but was adjusted on a subject-by-subject basis to correct for individuals’ abilities to perceive the stimulus over the noisy background in the MRI scanner. Like visual target events, “auditory-up” and “auditory-down” targets were marked by a linear increase or decrease in volume by 30% from this baseline over 1.6s and a linear return in the subsequent 0.8s, respectively.

In both experiments, participants first performed two mental counting blocks, during which they were asked to silently count target occurrences and report them to the experimenter after the end of the second “counting” block. In Experiment I, both counting blocks contained 30 “visual-down” targets. In Experiment II, one counting block contained 24 “visual-up” and the other 24 “visual-down” targets (Fig. 1B).

In both experiments, participants then performed eight experimental blocks that required them to press a button on a two-button MR-compatible response box with their right thumb as soon as they were certain that a target had occurred. In Experiment I, participants underwent two echo planar imaging (EPI) sessions that included all four conditions—auditory up, auditory down, visual up, and visual down. In Experiment II, each condition was recorded in its own EPI session (Fig. 1B). Again, each block contained 30 targets in Experiment I and 24 targets in Experiment II.

### 2.2 MRI Data acquisition

All MRI data were collected using a Philips 3T Achieva MRI scanner with a 32 channel head coil in Trinity College Institute of Neuroscience. For Experiment I, data were recorded in three EPI sessions containing 494 volumes lasting 16.5 min. Functional data were acquired using a T2*-weighted gradient echo, EPI sequence with the following parameters: 32 interleaved slices, 0.3 mm gap, 3.0 × 3.0 × 3.0 mm voxel size, 80 × 80 matrix size, 240 mm field of view (FOV), 25 ms echo time (TE), 2000 ms repetition time (TR), and 90° flip angle. Additionally, a high spatial resolution anatomical volume was obtained at the end of the session using a T1-weighted sequence with these parameters: 176 slices, 0.5 mm gap, 0.9 × 0.9 × 0.9 mm voxel size, 256 × 256 matrix size, 230 mm FOV, 2300 ms TR, 2.96 ms TE, and 9° flip angle. In between each EPI session, subjects had a two-minute rest whilst T1 and T2 weighted clinical scans were acquired. These images were not used for any analytical purposes.

In Experiment II, a high-resolution structural MRI was obtained using a T1-weighted sequence. This sequence featured 176 slices, 0.3 mm gap, 1.0×1.0×1.0 mm voxel size, 256 × 256 matrix size, 256 mm FOV, TE of 2.98 ms, TR of 2300.0 ms, and 9° flip angle. Functional imaging was carried out in six sessions using a T2*-weighted gradient EPI sequence. The functional sequence parameters were set at 36 slices, 0.5 mm gap, 3.0 × 3.0 × 3.0 mm voxel size, 64 × 64 matrix size, a 192 mm FOV, TE of 30.0 ms, TR of 2000 ms, and 80° flip angle.

### 2.3 EEG Data acquisition

For Experiment I, EEG data were collected simultaneously with the fMRI data using a 64-channel MR-compatible BrainCap system (Brain Products GmbH, Munich, Germany), digitized at 500 Hz. Of these channels, 63 electrodes were positioned on the participant’s scalp to acquire EEG and one electrode served as a monitor of the electrocardiogram (ECG). Event timing, participants’ responses and TTL pulse times were acquired as part of the EEG data set. For Experiment II, 128-channel EEG data were recorded at 512 Hz in separate experimental sessions outside of the MRI scanner. The locations of all electrodes were digitised relative to the Nasion, Left Pre-Auricular (LPA), and Right Pre-Auricular (RPA) points for each participant for matching to individual structural MRI in mean-field modelling (see below).

### 2.4 Data preprocessing

The fMRI data preprocessing procedure applied to the datasets includes slice-timing correction to adjust for differences in image acquisition times, high-pass filtering with a cutoff of 128 s to remove low-frequency noise, spatial smoothing using a Gaussian kernel of 6 mm to improve signal-to-noise ratio, and head motion correction to account for participants’ movements during the scan. Additionally, we used SPM12 software package^13^ for realignment of the fMRI images, extraction of motion parameters, co-registration of the mean fMRI volumes with the structural data for each participant, and normalisation to standard brain space.

Preprocessing of the EEG data followed a standardized pipeline designed to maximize signal quality while minimizing the influence of non-neural artifacts. Continuous EEG signals were first high-pass filtered at 0.1 Hz to remove slow drifts. To attenuate mains interference, narrow-band line noise centered at 50 Hz was suppressed across all channels. Channels exhibiting prolonged flatlining, excessive noise, or poor correlation with neighboring electrodes were automatically identified and subsequently interpolated using a spherical spline method based on the original sensor configuration. All data were re-referenced to the average of the remaining channels to enhance the spatial distribution of signals and to mitigate reference-related biases. Independent component analysis (ICA) was performed to decompose the signal into spatially fixed components, for the purpose of artifact correction using the extended Infomax algorithm implemented in EEGLAB (Delorme & Makeig, 2004). The resulting components were automatically classified based on their spatial topography and spectral content. To identify and remove artifactual independent components, we used the ICLabel (Pion-Tonachini et al., 2019). Components classified as non-brain (ocular, muscular, cardiac, channel noise, or line noise artifacts) with a probability greater than 0.5 were excluded from further analysis.

Following artifact correction, the signals were low-pass filtered at 80 Hz to reduce high-frequency noise and to preserve the spectral content relevant to event-related responses.

### 2.5 Behavioural analysis

For each participant and condition (auditory up/down, visual up/down), reaction times (RTs) from the fMRI sessions were compiled, and invalid responses (<0 or >5 s) were excluded. Kernel density estimation was then used to visualize condition-specific RT distributions.

### 2.6 BOLD-fMRI data analysis

We employed the Generalised Linear Model (GLM) technique within the SPM12 (Penny et al., 2011) fMRI toolbox to analyse our two fMRI datasets. This analysis included various experimental conditions to extract potential brain regions activated during detection decisions in a supramodal, motor-independent manner.

#### GLM framework and regressors

Our GLM analysis included session-specific regressors to account for variability between sessions, target regressors aligned with the target onsets in each session with 2.6 seconds duration, and 6 head motion regressors. All targets were modelled, regardless of response, using the other parts of each session as an implicit baseline for comparisons. This approach was uniformly applied to both datasets to maintain consistency in data analysis.

#### Hemodynamic response function (HRF)

We utilised a second-order Fourier basis function (5 regressors) with a 20-second Hanning window to model the BOLD response. This choice was aimed at capturing the complex dynamics of hemodynamic changes more accurately than the canonical HRF could allow (Balsters & Ramnani, 2008).

#### First-level analysis

We treated all conditions within the mental counting and motor-response tasks as independent from each other for each dataset. For T-contrast vector analysis, we assigned the same weight to all of the target regressors specific to each condition and zero to all other confounding regressors separately for each participant. We used linear Volterra expansion and a first-order autoregressive model for serial correlations without global normalisation.

#### Second-level analyses

For the second-level analysis, we pooled the T-contrasts of each condition from both datasets and conducted a conjunction analysis assuming six independent levels specific to our six conditions to extract brain regions showing common detection decision activity.

### 2.7 EEG data analysis

The pre-processed continuous EEG data were segmented into epochs surrounding experimental events of interest. Epoch boundaries were defined separately for stimulus-locked (response-locked) analyses, with a window ranging from 0.5 s (1.5 s) prior to event onset to 2.5 s (0.5 s) post-event.

To ensure data quality at the epoch level, a multi-criteria rejection scheme was applied. Trials with signal excursions exceeding ±100 μV in any channel were discarded. In addition, outlier detection was performed using probabilistic measures of joint signal distribution across sensors and trials. Epochs containing improbable activity patterns were excluded. Channels contributing excessively to poor-quality trials were flagged, and any epochs with more than four such channels were removed from further analysis. All channels marked as noisy during artifact screening were interpolated again following epoch rejection to ensure consistent topographic resolution across participants.

Each participant’s epochs were sorted by stimulus type and RT category, yielding condition-specific averages suitable for event-related potential (ERP) and topographic analyses.

#### 2.7.1 ERP analysis at CPP electrodes

To investigate the temporal characteristics of decision-related neural activity, we examined both stimulus-locked and response-locked ERPs at a predefined set of centroparietal electrodes where the CPP is typically focussed (O’Connell et al., 2012). Stimulus-locked ERPs were aligned to target onset, while response-locked ERPs were aligned to the participant’s button press. For each participant, trials were sorted into fast, medium, and slow response categories based on within-subject tertile splits of RT distributions. EEG signals were averaged across trials within each condition and RT tertile.

Group-level analyses were conducted by averaging these participant-level ERPs across all subjects in Experiment 2 and plotting the CPP waveforms as a function of task condition and decisional latency. Separate scalp averages were computed across the centroparietal region of interest and across all electrodes to allow both regional and global characterizations of the observed effects.

To characterise the effects of condition and RT on CPP scalar measures, we performed two-way repeated-measures ANOVAs with Condition (four levels: AU, AD, VU, VD) and RT bin (three levels: Fast, Medium, Slow) as within-subject factors, implemented in MATLAB. Five CPP scalar measures were examined: the stimulus-locked linear slope (0–1000 ms post-stimulus), the response-locked linear slope (−300 to 0 ms pre-response), the instantaneous (tangential) slope at 150 ms pre-response computed as the first derivative of the low-pass filtered (5 Hz) signal at that latency, the peak amplitude, and the half-peak latency, both measured over a 0–1500 ms stimulus-locked window. Where Mauchly’s test (Mauchly, 1940) indicated a violation of sphericity, the Greenhouse–Geisser correction (Greenhouse & Geisser, 1959) was applied to degrees of freedom and p-values. Bonferroni-corrected post-hoc pairwise comparisons (Bonferroni, 1936) were conducted on the main effects of both Condition and RT bin, as well as on RT bin effects within each condition separately. In addition, to obtain a single, clean summary statistic for the RT bin effect, a one-way repeated-measures ANOVA was performed on each scalar measure after collapsing data across conditions by averaging within each participant and RT bin.

To complement the ANOVA analyses, trial-level Pearson correlations (Pearson, 1900) were computed between each participant’s single-trial RT and the corresponding CPP scalar value. Fisher’s r-to-z transformation was applied to the individual correlation coefficients prior to group-level inference, and one-sample t-tests against zero were used to assess whether mean correlations differed significantly from chance across participants.

For all pairwise comparisons between adjacent sources in the ranked gain analysis (see below), normality of the paired differences was first assessed using the Lilliefors test (Lilliefors, 1967). Normally distributed differences were analysed with a paired t-test, with effect size reported as Cohen’s d (Cohen, 1988); non-normal differences were analysed with the Wilcoxon signed-rank test (Wilcoxon, 1945), with effect size reported as the rank-biserial correlation. A significance threshold of p < 0.05 was applied throughout, unless otherwise stated.

#### 2.7.2 Topographic mapping of response-locked EEG activity

To evaluate the spatial distribution of decision-related EEG signals, we computed scalp topographies at a fixed time point—50 ms prior to response onset. EEG signals were averaged across trials within each participant and condition, and subsequently across participants. We produced separate maps for each of the four target types and three RT bins, as well as a composite map that averaged over all conditions, RTs and participants. To identify the most informative recording sites, we computed the average amplitude at each electrode and identified the five electrodes with the highest mean pre-response amplitude, to be used in subsequent initial source contribution analyses.

#### 2.7.3 Forward modeling of EEG from fMRI-informed sources

Forward models were constructed using the FieldTrip toolbox (Oostenveld et al., 2011), incorporating individual anatomical MRI data, digitized EEG electrode positions, and source coordinates derived from fMRI. This procedure was applied to data from the second experiment only, where EEG and fMRI were recorded in separate sessions, free of MR-induced artifacts and with individual electrode digitization available.

For each participant, fiducial landmarks—nasion and bilateral preauricular points— were manually extracted from their structural MRI scans and used to realign the head volume in native space. The anatomical images were subsequently warped to MNI space using a nonlinear registration procedure to facilitate the projection of group-level source coordinates into individual head space.

The FEM approach (Wolters et al., 2006) was used to model the head volume. The T1-weighted anatomical images were segmented into five tissue compartments: scalp, skull, cerebrospinal fluid (CSF), grey matter, and white matter. Each compartment was discretized into a hexahedral mesh, and standard conductivity values were assigned to each tissue class. The head model was constructed from this mesh, allowing accurate estimation of the leadfield matrix that maps dipole sources to EEG scalp sensors.

EEG electrodes were realigned from the participant-specific digitized positions to the realigned MRI in MNI space. This was achieved by aligning the fiducials recorded during EEG electrode navigation to the fiducials defined in MNI coordinates, followed by warping and projection onto the scalp surface. All co-registration steps were performed in FieldTrip.

We generated the source model from the locations corresponding to the centers of significant clusters identified from the fMRI conjunction analysis and subsequently computed the individualized leadfield matrices. These matrices quantified the linear mapping from dipolar source activity to EEG sensor space, forming the basis for all forward modeling analyses.

#### 2.7.4 Single Source Contribution Analysis

In a preliminary step (reported in Supplementary Materials), to estimate the influence of each fMRI-informed source on the centroparietal EEG signal, we conducted a source-wise analysis of leadfield contributions across participants. Using each individual’s forward model, we computed the scalp projection pattern for every dipolar source and quantified its contribution to the centroparietal region of interest. Contributions were calculated based on the absolute sum of forward model vectors across the five centroparietal electrodes with the highest average amplitude at −50 ms, separately identified for each stimulus type.

For each participant, we extracted the forward model values associated with all dipoles and evaluated their influence on each region-of-interest (ROI) electrode. The contributions of each source were averaged across ROI electrodes and stored for statistical comparison. This allowed us to build a source × electrode contribution matrix for each subject and stimulus condition.

To identify sources with significantly stronger contributions, we applied a non-parametric Kruskal–Wallis test (Kruskal & Wallis, 1952) across sources for each electrode. The most influential source per electrode was identified based on mean contribution magnitude, and its dominance was evaluated through post hoc comparisons with all other sources using Bonferroni-corrected pairwise tests (Bonferroni, 1936).

#### 2.7.5 Pairwise Source Contribution Analysis

To investigate whether bilateral or functionally coupled regions exhibit synergistic influence on centroparietal scalp activity, we conducted a pairwise source contribution analysis. Source pairs were defined based on anatomical similarity and hemispheric symmetry, pairing homologous or closely adjacent regions. For each participant and stimulus condition, forward model projections were computed for each pair by summing the respective leadfield vectors and estimating their combined influence at centroparietal electrodes.

For each source pair, the magnitude of the combined scalp projection was computed as the Euclidean norm of the summed forward model vectors across the three orthogonal dipole components. Contributions were then extracted for the five centroparietal electrodes previously identified as exhibiting maximal CPP activity. This process yielded a contribution matrix representing the influence of each pair across electrodes, participants, and stimulus conditions.

To determine which source pairs contributed significantly more than others, we performed non-parametric Kruskal–Wallis tests across all defined pairs for each ROI electrode. The pair with the highest mean contribution was identified, and Bonferroni-corrected post hoc comparisons were used to assess statistically significant differences with all other pairs. These analyses were repeated independently for each stimulus type.

#### 2.7.6 Forward projection of fMRI-informed source activity to EEG topography

To directly assess whether fMRI-identified cortical sources could account for the scalp distribution of decision-related EEG signals, we implemented a forward modeling pipeline that estimated scalp potentials from the spatial configuration and strength of dipolar sources. The approach integrated fMRI contrast data, individualized leadfield models, and group-level EEG topographies.

Source locations were defined as the peak coordinates of activation clusters obtained from a group-level conjunction analysis across task conditions in the fMRI data. For each source, its corresponding dipole strength was extracted by sampling the unthresholded t-statistic value at that coordinate from the statistical parametric map.

To define orientation vectors for each dipole, we used a participant-specific finite element head model to estimate the local cortical surface normal. For each source, orientations were computed for all participants by averaging polygon normals within a growing neighborhood around the MNI location until a stable, non-zero direction was identified. The final orientation vector per source was computed as the mean of participant-level directions.

EEG topographies at −100 to 50 ms relative to response onset with 25 ms step were computed by averaging response-locked ERP data across all conditions and participants. Using these empirical topographies as a target, we projected the modeled dipole activity to the scalp by computing the forward projection of each source (using the average orientation and dipole strength) through the individualized leadfield matrices, and averaging the resulting sensor-level projections (predictors) across participants.

To mitigate multicollinearity among the fMRI-informed predictors in our EEG forward model, we applied a clustering-based dimensionality reduction step prior to estimating gain contributions. First, we projected all dipolar sources—each weighted by their estimated orientation and fMRI-derived strength—onto the EEG sensor space using participant-specific leadfields. This resulted in a matrix of predicted source topographies. We computed the pairwise Pearson correlation matrix across all projected source topographies and converted it into a distance matrix using the absolute correlation (distance = 1 − |correlation|). Hierarchical clustering (average linkage) was then applied to group highly correlated sources. To ensure numerical stability, we selected the correlation cut-off to yield a reduced design matrix with a condition number below 30, in line with known guidelines (David A. Belsley et al., 1980).

For each resulting cluster, the topographies of its members were averaged to form a single predictor. Gain estimation was then performed via least-squares fitting of these reduced predictors to the observed response-locked EEG. The resulting gains represented each cluster’s contribution to the measured scalp signal. Columns of the reduced design matrix were z-scored before computing the model’s condition number.

To evaluate the strength and reliability of each cluster’s contribution, we computed the product of the fitted gain and dipole strength for each cluster and normalized these values across all sources and participants. Clusters were then ranked by their mean contribution. For adjacent clusters in this ranking, we compared the distributions of their time-resolved gain values. Prior to statistical testing, the normality of the paired differences was assessed (Lilliefors test). If the differences were normally distributed, a paired *t*-test was used; otherwise, the Wilcoxon signed-rank test (Wilcoxon, 1945) was applied (*p* < 0.05). This procedure accounts for potential violations of normality while preserving the paired structure of source contributions across time.

#### Leave-one-cluster-out ablation analysis

To evaluate whether ranking clusters by fitted gain magnitude reflects each cluster’s true, unique contribution to model fit — rather than partly reflecting shared variance with correlated neighboring clusters — we performed a leave-one-cluster-out (LOO) ablation analysis. For the full model (all the clusters), gain was estimated via ordinary least squares at each of the seven response-locked timepoints analysed (−100 to +50 ms in 25 ms steps), and the resulting reconstructed topography was compared against the observed topography to obtain a normalized MSE at each timepoint (mean squared error divided by the variance of the observed data at that timepoint). This procedure was then repeated for the number of clusters, each time excluding exactly one cluster’s predictor column from the design matrix and refitting the remaining clusters via the same least-squares procedure. For each excluded cluster and each timepoint, we computed ΔMSE as the difference between the MSE of the reduced (N−1 cluster) model and the MSE of the full model; a larger ΔMSE indicates that removing that cluster degraded model fit more, and therefore that the cluster made a larger unique contribution not redundant with the remaining clusters. Clusters were ranked by mean ΔMSE across the seven timepoints. To formally compare adjacent-ranked clusters, paired differences in ΔMSE across the seven timepoints were tested for normality (Lilliefors test) and compared using a paired t-test (if normally distributed) or Wilcoxon signed-rank test (if not). Agreement between the gain-based and ablation-based rankings was assessed using Spearman’s rank correlation coefficient across all the clusters.

#### Model fit as a function of source count

To assess how many clusters are required to reconstruct the observed scalp topography, and whether this depended on how clusters were prioritized, we refit the forward model using only the top-K clusters, separately ordering clusters by gain magnitude and by ablation importance (as defined above). For each K and ranking, gain was re-estimated via ordinary least squares using only the top-K clusters’ predictor columns, separately at each of the seven response-locked timepoints analyzed (−100 to +50 ms, 25 ms steps), and the resulting reconstructed topography was compared against the observed topography to obtain a normalized MSE at each timepoint, which was then averaged across timepoints to give one value per K. To visualize this progression spatially for K = 1–5, we additionally reconstructed the time-averaged (across the seven timepoints) scalp topography using only the top-K clusters under each ranking; the observed time-averaged topography is shown once, since it does not depend on ranking or K. For the corresponding electrode-wise fit error, we computed, for each channel and each K, the mean squared difference between the top-K reconstructed and observed signal across the seven timepoints, normalized by that channel’s own observed variance across those timepoints — the same normalization convention used for the overall MSE, computed per channel instead of per timepoint.

#### Mental-counting control analysis

To test whether the cortical contributions identified for the response-locked CPP depend on manual response, we refitted the same forward model to the CPP recorded in the mental-counting blocks (CU and CD). Stimulus-locked epochs (−500 to 2500 ms relative to target onset) were averaged separately for each participant and counting condition. Each average was then low-pass filtered (fourth-order Butterworth, 10 Hz cut-off, zero-phase) to remove the steady-state response to the 21.25-Hz annulus flicker, which is phase-locked to target onset and therefore survives trial averaging.

Counting trials have no response event against which single trials can be aligned. The analysis window was therefore anchored to the peak of the grand-average CPP (ROI electrodes A2, A3, A4, B2 and D16), pooled over CD and CU and searched between 200 and 2200 ms after target onset. Scalp topographies were extracted at seven timepoints from 100 ms before to 50 ms after this peak in 25-ms steps, matching the seven response-locked timepoints (−100 to +50 ms) of the button-press analysis. For a like-for-like comparison, the button-press topographies were re-extracted with the identical 10-Hz filter.

Both conditions were fitted with the same predictor matrix as the main analysis. At each timepoint, cluster gains were estimated by ordinary least squares. Each cluster’s unique contribution was quantified by leave-one-cluster-out ablation: ΔMSE, the increase in normalised mean squared error when that cluster is removed and the model is refitted. Within each condition, clusters were ranked by their mean |gain| and by their mean ΔMSE across the seven timepoints. Agreement between the button-press and mental-counting rankings was quantified with Spearman’s rank correlation across the clusters, and clusters ranked in the top five in both conditions were taken as shared contributors.

#### Participant-level forward-model fits

To test whether the group-average rankings held across individuals, we refitted the forward model separately for each participant. Each participant’s design matrix was built from their own FEM leadfield. It used the same conjunction peaks and surface-normal orientations as the group model, restricted to the channels shared by that participant’s EEG and leadfield, and z-scored across channels. Observed topographies were taken from each participant’s own average. The windows and filter matched the group analysis: seven timepoints around the response for button press, and seven timepoints around the pooled group CPP peak for mental counting, with a 10-Hz zero-phase low-pass filter in both conditions.

Individual designs are more ill-conditioned than the group-average design. We therefore raised the clustering threshold until every participant’s clustered design had a condition number below 30. Cluster membership was defined on the group model and applied to every participant. Each cluster predictor was the mean of that participant’s member topographies, re-standardised to unit variance. Gains and leave-one-cluster-out ΔMSE were estimated at each timepoint and averaged over the seven timepoints.

For each cluster, the participants’ mean signed gains were tested against zero with a sign-flip permutation test (10,000 permutations). Family-wise error across clusters was controlled with the max-t statistic. To give each cluster’s group rank a 95% interval, we recomputed the rankings by mean |gain| and mean ΔMSE in 10,000 bootstrap resamples of participants. Because in-sample ΔMSE is non-negative by construction, unique contributions were also assessed out of sample: gains were estimated on N−1 participants (leave one subject out) and ΔMSE was evaluated on the held-out participant. Agreement between the two conditions’ rankings was quantified with Spearman’s ρ, with bootstrap 95% CIs. Condition differences in each cluster’s share of total ΔMSE were tested with a paired permutation test, using the participants who contributed to both conditions.

### 2.8 Software and hardware

We employed the EEGLAB toolbox (Delorme & Makeig, 2004) for preprocessing EEG data and utilised the SPM, version 12 (SPM12) software package in MATLAB (Version 2026b) for analysing fMRI data, and statistical analyses. We implemented the EEG source analysis in the FieldTrip toolbox (Oostenveld et al., 2011). All fMRI analyses as well as ERP preprocessing were conducted on the Northern Ireland Performance Computing (NI-HPC) facility using the Kelvin2 system (www.ni-hpc.ac.uk). Source code used in our analysis will be made available upon publication.

## 3 Results

In both experiments, participants engaged in a continuous target detection task across five blocked conditions involving flickering visual and auditory stimuli (Fig. 1), with targets defined by a linear ramp-up or ramp-down in visual contrast or auditory volume, randomly separated by inter-target intervals (ITIs) of 4, 6, or 8 s (Fig. 1A). Targets thus gradually emerged but were ultimately easy to detect, leading to high hit rates (94.5% ± 9.0) and long and variable RTs (Fig. 2A).

**Figure 2.**
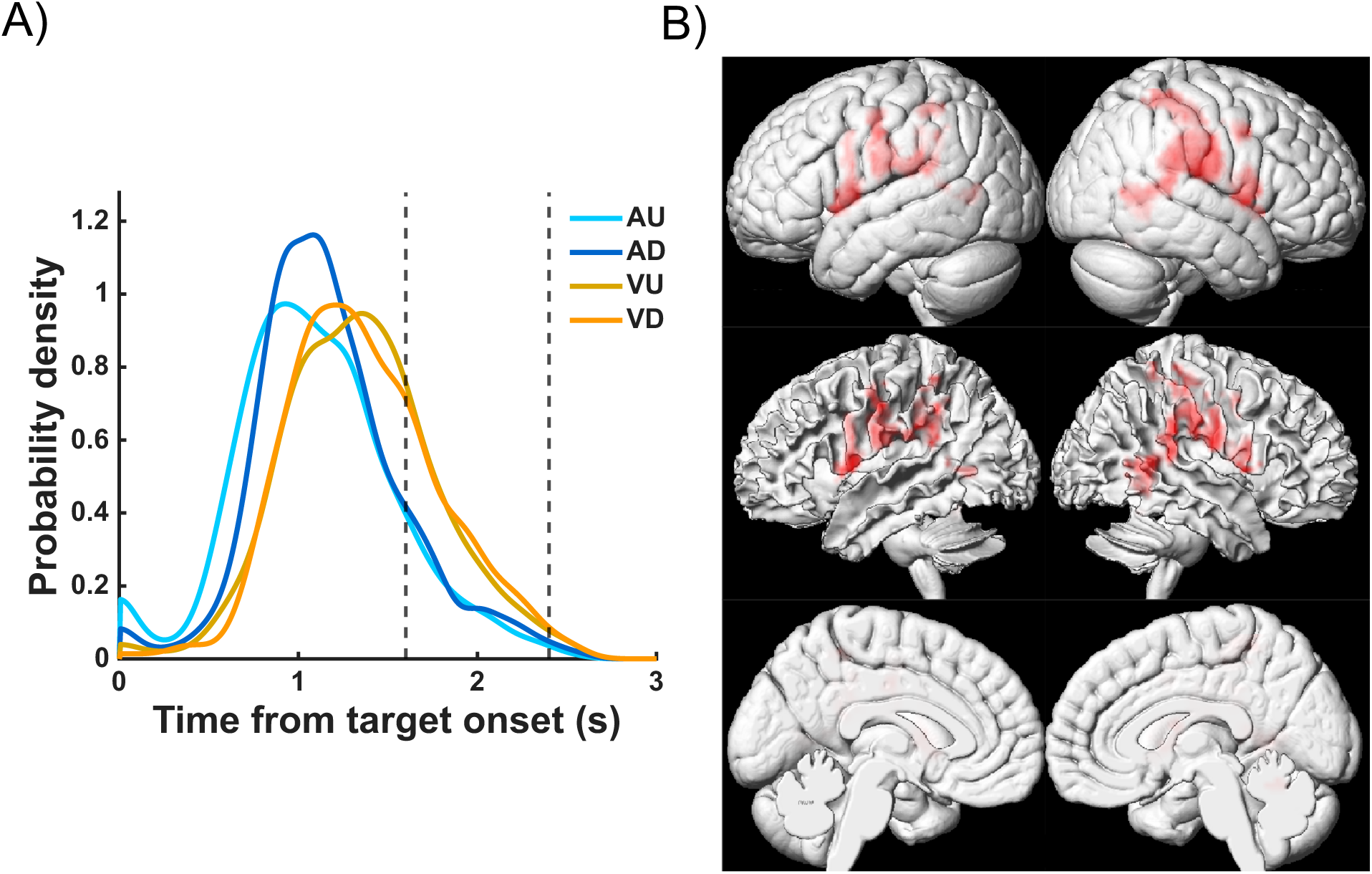
Potential supramodal decision-related regions in the parietal and frontal lobes as identified by conjunction analysis. A) Aggregated reaction time (RT) distributions of auditory up (AU), auditory down (AD), visual up (VU), and visual down (VD) conditions, pooled from fMRI sessions of Experiments I and II. B) The identified regions with decision-dependent activity are the postcentral gyrus, supramarginal gyrus (inferior parietal lobule), and superior parietal lobule (SPL) in the parietal cortex as well as the precentral gyrus (PRG), central opercular cortex, and middle frontal gyrus.

In the “mental counting” condition, participants counted targets during an experimental block and reported the number of detected targets back to the experimenter verbally after the end of the block (Fig. 1B). In contrast, in motor response blocks, participants responded by pressing a button on a two-button MR-compatible response box using their right thumb, recording choices and precise RTs. Across the 45 participants (31 from the separate fMRI dataset and 14 from the simultaneous EEG-fMRI dataset), auditory targets were detected significantly faster than visual targets, with mean RTs of 1.1 ± 0.4 s for auditory up/down compared to 1.3 ± 0.4 s for visual up/down (all p < 0.001, paired t-tests).

### 3.1 Conjunction analysis identifies common, supramodal decision-related regions across tasks

We analyzed blood oxygenation level-dependent (BOLD) fMRI data using the generalised linear model (GLM) framework implemented in SPM12 (Penny et al., 2011). T-contrasts from both datasets were pooled and subjected to a conjunction analysis to identify brain regions engaged across all experimental conditions (p < 0.05, uncorrected, Fig. 2B, Tables S1 and S2). Since these conditions vary in the sensory and motor-related regions that they activate, the identified regions should include brain areas that encode abstract perceptual decision processes. Several of the identified sources share the same coarse anatomical label because the fMRI conjunction analysis identified multiple distinct local maxima within a small number of large clusters. Our analysis identified the postcentral gyrus, supramarginal gyrus (SMG, part of inferior parietal lobule (IPL)), and superior parietal lobule (SPL) in the parietal cortex as well as the precentral gyrus (PRG), central opercular cortex, and middle frontal gyrus in the frontal cortex (Fig. 2B). Interestingly, the identified regions additionally included a part of the cerebellum, a structure traditionally more associated with motor control but increasingly implicated in higher cognitive functions including perceptual decision-making and evidence evaluation (Schmahmann, 2019).

### 3.2 EEG signatures of abstract evidence accumulation

To assess how decision-related activity unfolded over time in this dataset, we examined both stimulus-locked and response-locked EEG data at centroparietal scalp sites across visual and auditory conditions. In line with previous findings, the CPP had a centroparietal focus and exhibited a gradual build-up after target onset that peaked around the average response time (Fig. 3A, see Fig. S1A for mental counting and Fig. S1B for trial-by-trial response). When waveforms were collapsed across conditions and plotted separately as a function of RT bin (Fig. 3A bottom), the CPP’s peak latency varied in lockstep with RT, with a similar amplitude reached just before the response regardless of RT, consistent with the purported role of the CPP in determining decision termination time through reaching a criterion amount of cumulative evidence (O’Connell et al., 2012). Although peak CPP amplitude varied across conditions (two-way RM-ANOVA: F (3,93) = 5.45, p = 0.002; VU larger than AD (p = 0.011) and AU (p = 0.041)), topographies for all conditions included a clear centroparietal focus and waveforms similarly showed earlier buildup for faster RTs and a peak and fall shortly following RT (Fig. 3B and Fig. S2).

**Figure 3.**
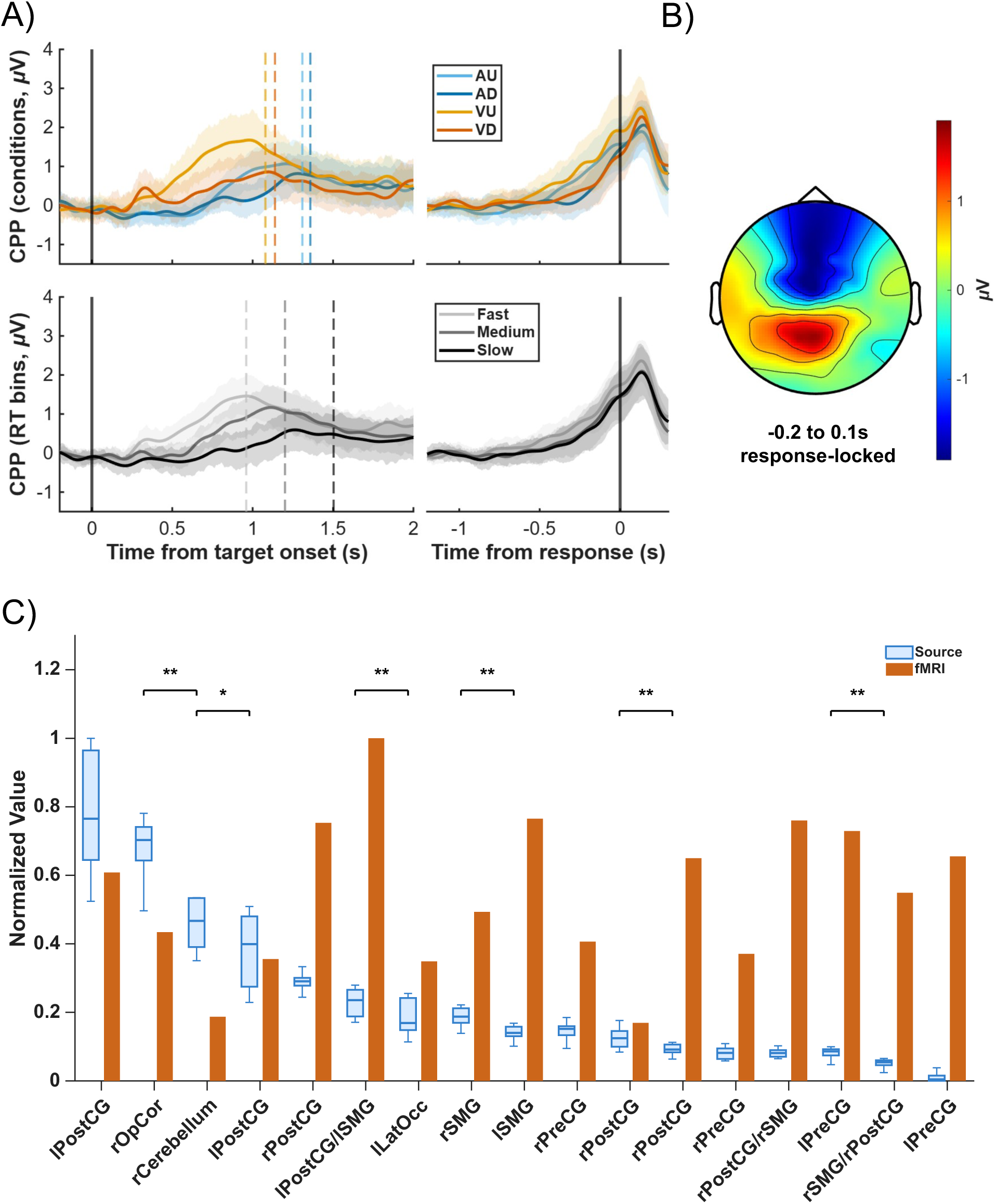
Decision-related brain sources contribute to the scalp decision signal. A) Grand average CPP waveforms in different conditions and RT bins during the button press task. B) Corresponding topographic map of response-locked EEG activity averaged from −200 ms to 100 ms from button press. C) Ranked, normalized contributions of the conjunctively-activated fMRI-sources to the response-locked CPP topography. Boxplots show gain distributions estimated for each 25 ms step from -100 to 50 ms response-locked (see Methods for details). Where multiple sources share the same coarse anatomical label (e.g., multiple peaks within left postcentral gyrus), each corresponds to a spatially distinct local maximum from the fMRI conjunction analysis—see Table S2 for exact peak coordinates and parent cluster assignments. Bars show fMRI conjunction activations at the same sources. Significant pairwise differences between source contributions marked with stars (\**p* < 0.05, \*\**p* < 0.01), with line colors denoting effect size (black = large). Source abbreviations: lPostCG = left postcentral gyrus; rOpCor = right opercular cortex; rCerebellum = right cerebellum; rPostCG = right postcentral gyrus; lPostCG/lSMG = left postcentral gyrus / left supramarginal gyrus; lLatOcc = left lateral occipital cortex; rSMG = right supramarginal gyrus; lSMG = left supramarginal gyrus; rPreCG = right precentral gyrus; rPostCG/rSMG = right postcentral gyrus / right supramarginal gyrus; lPreCG = left precentral gyrus; rSMG/rPostCG = right supramarginal gyrus / right postcentral gyrus. Prefix l = left hemisphere, r = right hemisphere.

Aside from this common centroparietal positive focus, the two auditory conditions (AU and AD) displayed a prominent frontocentral negativity that was absent in the visual conditions (Fig. S2B), most likely reflecting superimposed auditory sensory responses from auditory cortex that are co-occurring with the decision-related centroparietal activity. The visual conditions, in turn, showed additional negativity over occipital sites, consistent with the expected contribution of visual cortical responses to the scalp topography. These condition-specific topographic features are to be expected given the different sensory systems engaged by auditory and visual stimulation, and they overlay the common centroparietal decision signal shared across conditions.

Together, these results confirm that a central CPP is a reliable feature of decision formation across modalities and response times and that the topographic differences between conditions principally reflect the superposition of modality-specific sensory components rather than fundamental differences in the decision accumulation process itself.

### 3.3 Cluster-based fMRI-informed forward modeling identifies parietal-opercular generators

Finally, after analysing single and pairwise source contributions (see Methods and Figs. S3-S10), we tested whether the full response-locked EEG topography could be reconstructed from fMRI-identified sources. We modeled EEG activity from −100 to +50 ms in each of seven 25-ms steps relative to response using 24 dipolar sources extracted from the fMRI analysis (Fig. S11) through clustered source projections to reduce collinearity among regressors, which resulted in using 17 clusters (Table S3). Fitting all cluster-based topographies to the observed EEG in one regression using least-square error estimation yielded a normalized mean squared error (MSE) of 0.32, indicating a strong correspondence between the fMRI-informed projections and the empirical CPP topography. The forward-projected topography predictors were each z-scored, so that the sources’ relative contribution or “gain” to the observed topography could be estimated directly from the standardized regression coefficients. The largest gain values were associated with a cluster spanning the left postcentral gyrus, followed by clusters in the right opercular cortex and right cerebellum (Fig. 3C, Fig. S12 and Table S3). While contributions from left postcentral gyrus and right opercular clusters did not differ significantly, both showed substantially and significantly greater explanatory power than lower-ranked sources including the right cerebellum.

To assess whether gain magnitude reflects each cluster’s unique contribution to the model fit, we performed a leave-one-cluster-out ablation analysis, refitting the model with each cluster excluded in turn and computing the resulting increase in MSE (ΔMSE) at each analyzed timepoint (Fig. S13, Table S4). Gain-based and ablation-based rankings were positively correlated overall (Spearman’s ρ = 0.87), and four of the top five clusters were shared between the two rankings (Tables S3–S4), supporting gain magnitude as a reasonable, though imperfect, proxy for a cluster’s unique contribution to the model fit.

To directly test how many sources are needed to reconstruct the observed topography, and whether this differed between gain-based and ablation-based rankings, we refit the model using only the top-K clusters (K = 1 to 17) added in each ranking’s order (Fig. 4A). Normalized MSE decreased sharply and similarly under both rankings, reaching within 5% of the full-model MSE by K ≈ 10–11 clusters, indicating that a subset of top-ranked clusters accounts for most of the model’s explainable fit regardless of which ranking is used to select them. Reconstructing the topography from only the top-ranked clusters (K = 1–5) selectively and progressively recovered the centroparietal positivity characteristic of the CPP as K increased, under both rankings, while the corresponding electrode-wise error maps showed this improvement was concentrated over centroparietal electrodes rather than distributed evenly across the scalp (Fig. 4B–C). This convergence toward the observed CPP topography using only a handful of fMRI-derived clusters — the same clusters implicated by the conjunction analysis — supports the interpretation that these extracted regions are plausible neural generators of the CPP, rather than an incidental byproduct of the whole-brain model fit.

**Figure 4.**
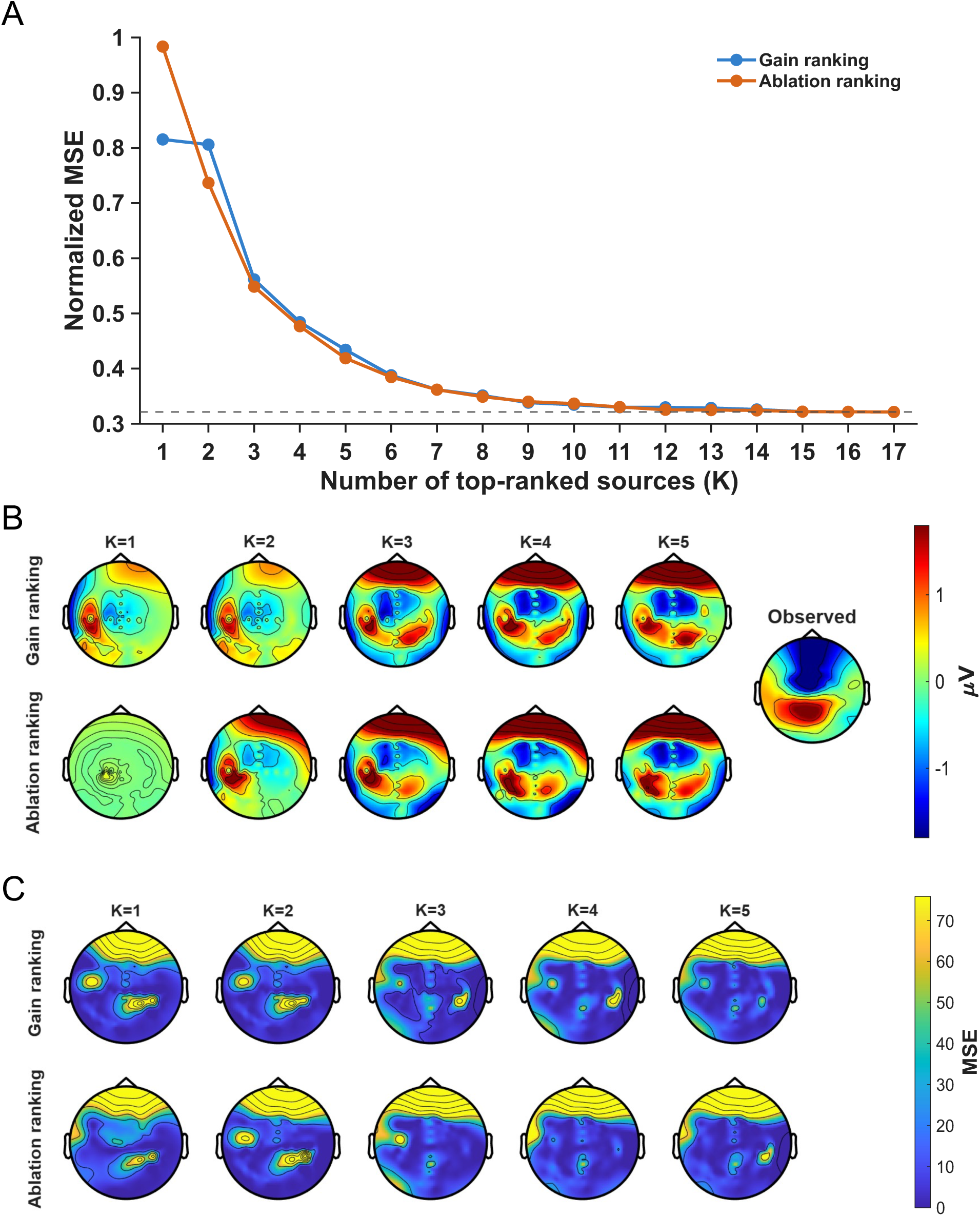
The fMRI-conjunction-derived sources selectively and increasingly explain the CPP-relevant centroparietal topography as more sources are included, without a corresponding improvement at frontal electrodes. (A) Normalized mean squared error (MSE) of the reconstructed scalp topography as a function of the number of top-ranked clusters (K) included in the model, for K = 1 to 17 (all clusters). Clusters were added in order of either gain magnitude (blue) or ablation importance (ΔMSE upon leave-one-cluster-out removal; orange). Both curves converge to the same value at K = 17, since the full model is identical regardless of the order clusters were added in. Dashed horizontal line indicates the full-model (K = 17) MSE, provided as a reference for how quickly each ranking approaches asymptotic fit quality. (B) Reconstructed scalp potential (µV) topographies for K = 1–5, using only the top-K clusters by gain ranking (top row) or ablation ranking (bottom row), time-averaged across the analyzed response-locked window (−100 to +50 ms, 25 ms steps). The rightmost panel shows the observed topography for comparison (identical for both rows, shown once). All panels share one colour scale. (C) Electrode-wise fit error corresponding to panel B: for each channel, mean squared error between the top-K fitted and observed signal across the analyzed time window, normalized by that channel’s own observed variance (unitless), for K = 1–5 under gain ranking (top row) and ablation ranking (bottom row). All panels share one colour scale, separate from panel B. The gain and ablation rankings assess related but non-equivalent aspects of source contribution. Gain measures the fitted weighting of each cluster in the joint reconstruction, whereas ablation ΔMSE measures the non-redundant loss of fit produced when that cluster is removed and the remaining clusters are refitted. The high correspondence between rankings therefore indicates that the dominant source subset is relatively robust to whether importance is defined by fitted coefficient magnitude or by loss of model fit. Notably, the Top-5 reconstructions improved the spatial match primarily over the centroparietal region containing the CPP, while comparatively large residual errors persisted over frontal electrodes. Because whole-scalp MSE continued to improve beyond five clusters, this result does not imply that the Top-5 explain the complete scalp topography; rather, it indicates that a small subset of the fMRI-derived sources preferentially accounts for the CPP-relevant component of that topography.

### 3.4 Parietal-opercular regions contribute to the CPP regardless of motor involvement

To test whether the source contributions to the CPP depended on manual response, we refitted the forward model to the CPP recorded during mental counting. Both conditions were fitted with the identical 17-cluster, strength-free forward model, and clusters were ranked within each condition by mean absolute gain and by leave-one-cluster-out ablation (ΔMSE) across the seven timepoints (see Methods).

In the absence of any manual response, the grand-average CPP rose from approximately 400 ms after target onset and peaked at 869 ms (Fig. S14A). At its peak, it still showed a centroparietal positivity with a posterior negativity, a topography that persisted for several hundred milliseconds (Fig. S14B). Remarkably, the forward model accounted for the mental-counting topography nearly as well as for the button-press topography (normalised MSE = 0.39 and 0.32, respectively). Across all 17 clusters, rank agreement between conditions was modest for both gain (Spearman ρ = 0.24, p = .35) and ablation (ρ = 0.24, p = .36).

Three of the five clusters with the largest gains were common to both conditions (Fig. 5A): the left postcentral cluster, the right opercular cortex and the right postcentral gyrus. Ranked by unique contribution, two clusters were among the five strongest in both conditions (Fig. 5B): the right opercular cortex and the left lateral occipital cortex. The right opercular cortex was therefore the only cluster ranked among the five strongest contributors in both conditions on both measures.

**Figure 5.**
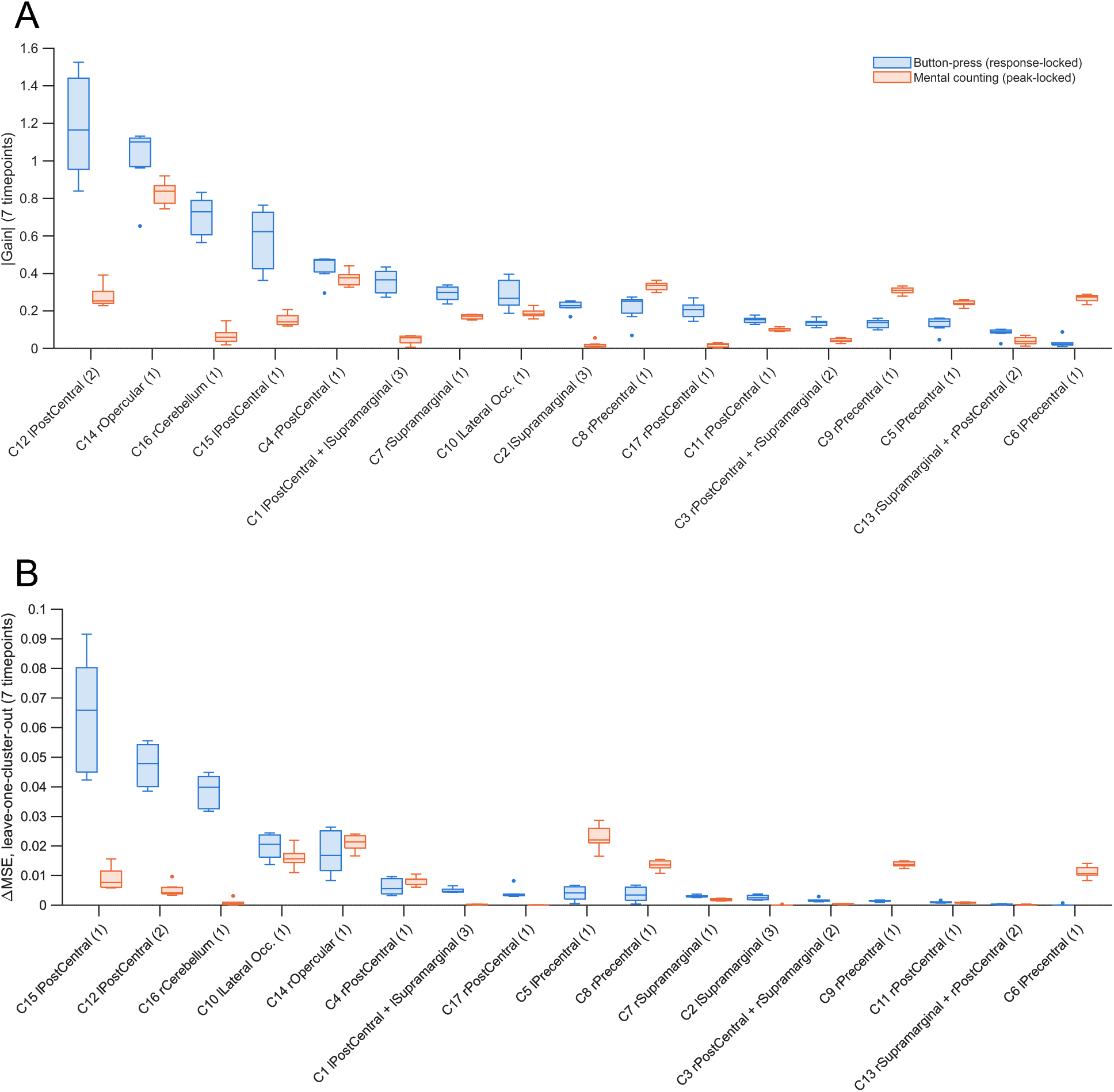
Cluster contributions to the button-press and mental-counting CPP (group-average fit). (A) Absolute gain (see Table S3 for MNI coordinates) and (B) leave-one-cluster-out ablation (ΔMSE) (see Table S4 for MNI coordinates) for the 17 source clusters. Each box summarises the seven analysed timepoints. Clusters are ordered by their button-press rank; the number in brackets is the number of member sources. Both conditions were fitted with the same forward model, so they differ only in the observed topography. The full-model normalised MSE was 0.32 for button press and 0.39 for mental counting. The right opercular cortex (48, 2, 5) ranked second (button press) and first (mental counting) by gain, and fifth and second by ablation. It was the only cluster among the five strongest contributors in both conditions on both measures. Without a manual response, the right cerebellum (24, −58, −19) fell from third to 12th by gain and from third to 11th by ablation. The left postcentral/superior parietal source (−48, −40, 56) fell from fourth to tenth by gain and from first to seventh by ablation. Spearman ρ gives the rank agreement between conditions across the 17 clusters (gain ρ = 0.24, p = .35; ablation ρ = 0.24, p = .36).

We next refitted the model separately for each participant (button press, N = 31; mental counting, N = 29), using 10 clusters so that every individual design was well conditioned (condition number < 30). At this threshold the right opercular cortex merges with the adjacent right postcentral source, which also ranked among the top five in the group analysis. This merged cluster was the only cluster ranked among the five strongest contributors in both conditions on both measures (Fig. S15 and Table S5). The clustered model preserved the main features of the grand-mean topographies in both conditions (Fig. S16): the centroparietal positivity and the frontal negativity for button press, and the centroparietal positivity with a posterior negativity for mental counting.

Together, these findings support the hypothesis that the CPP is primarily generated by bilateral parietal-opercular regions, with a dominant contribution from right opercular cortex.

## 4 Discussion

The present study sets out to identify brain regions that support abstract perceptual decision-making, i.e. neural processes that are active independently of the sensory modality of the stimulus and the motor requirements of the task. To address this question, we combined BOLD-fMRI and EEG data through a multimodal source analysis framework. Using BOLD-fMRI data, we first applied a conjunction contrast analysis across auditory and visual modalities as well as across different motor requirements (mental counting and button press). This revealed a bilateral cortical network including the postcentral gyrus, supramarginal gyrus (SMG, a part of the inferior parietal lobule, IPL), superior parietal lobule (SPL), precentral gyrus, central operculum, middle frontal gyrus, and additionally the cerebellum.

In parallel, EEG recordings showed response-locked activity over central electrodes resembling the well-studied CPP activity, which ramped up over time toward decision commitment. We then used the fMRI-defined clusters to inform the forward model of EEG source activity, i.e., the centers of the clusters were taken as priors for estimating the leadfield using finite element method (FEM) for head modeling. By projecting these sources to the scalp and quantifying their contributions to the observed CPP-like activity in the –100 to +50 ms with respect to the motor response window, we identified the parietal–opercular regions as the most dominant contributors. In comparison with previous studies on CPP activity, our multimodal approach adds spatial specificity. Whereas EEG alone can tell when the decision variable peaked, combining EEG with fMRI (and FEM of head modeling) allows us to suggest where in the brain this buildup was occurring.

### 4.1 Distributed frontoparietal network supports abstract perceptual decision-making

Our fMRI conjunction analysis points to a broad frontoparietal network that consistently underpins decision formation regardless of sensory modality. This aligns well with prior research in both animals and humans. In non-human primates, neurons in the parietal cortex – classically area LIP on the lateral bank of the intraparietal sulcus (IPS) – ramp up their firing rates as evidence is accumulated for a decision (Lafuente et al., 2015; Seideman et al., 2022; Zylberberg & Shadlen, 2025). Similar “accumulation-to-bound” dynamics have been observed in rodent decision-making circuits: neurons in rodent posterior parietal cortex (PPC) and frontal orienting fields (an anterior motor-planning region, leveraging accumulated evidence for decisions) also exhibit gradual changes in activity reflecting evidence accumulation (Boyd-Meredith et al., 2022; Erlich et al., 2015; Gupta et al., 2024). However, previous human fMRI work on abstract decision-making has often emphasized more circumscribed regions—particularly the IPS, superior parietal cortex, or dorsolateral prefrontal cortex (DLPFC)—as hubs for supramodal decision signals (e.g., Morito & Murata (2022) and Pisauro et al. (2017)).

Crucially, our finding that bilateral SPL and postcentral gyri are engaged by decisions across modalities dovetails with human studies of “domain-general” decision signals. The SMG (part of IPL) is thought to integrate multisensory information and has been linked to updating decisions or switching strategies (Igelström & Graziano, 2017), and it borders the IPS where neurons encode accumulated evidence in primates (Gold & Shadlen, 2007). The SPL, on the dorsal aspect of parietal cortex, plays a role in spatial attention and sensorimotor integration; its activation during decisions likely reflects the allocation of attention to accumulating sensory evidence and the transformation of that evidence into a motor plan (Rushworth & Taylor, 2006). Indeed, non-human primate studies indicate parallel decision pathways in the parietal cortex for different effectors (e.g. reach-related SPL vs. saccade-related LIP), both carrying an evolving decision variable (Lafuente et al., 2015). The activation of the postcentral gyrus, mainly known as the site of the primary somatosensory cortex, is unexpected; although its activation could reflect somatosensory attention or the processing of sensory feedback related to the decision (Eickhoff et al., 2008), it is not yet clear why this would occur in this task, and will bear following up in future research.

Our results show middle frontal activation across modalities, consistent with the view that this region provides supramodal support for decision formation, especially when decisions require maintaining a rule (Miller & Cohen, 2001) or comparing evidence against a criterion (Heekeren et al., 2004; Philiastides et al., 2011).

Finally, highlighted by our fMRI conjunction, the opercular regions (central operculum) likely correspond to the frontoparietal opercular cortex overlying the insula. These opercular areas are part of the so-called cingulo-opercular network, which is associated with sustaining task sets and monitoring performance (Dosenbach et al., 2008). In decision-making contexts, the cingulo-opercular network is thought to signal the need for increased cognitive control – for instance, when evidence is conflicting or a decision is especially difficult (Pierce et al., 2023). Thus, the opercular activation in our experimental task may relate to the engagement of this control network to support accurate performance across different types of decisions.

It is noteworthy that in some rapid or highly practiced decisions, the brain can minimize involvement of anterior frontal regions, relying more on parietal sensorimotor circuits (Tosoni et al., 2008).

### 4.2 Left parietal and right opercular cortical activities as candidate sources for CPP activity

By leveraging our fMRI results to constrain EEG source modeling, we obtained evidence that the CPP’s dominant generators lie in specific cortical areas – notably the left inferior parietal cortex (near the supramarginal/postcentral region) and right frontal or parietal opercular region (encompassing central operculum and possibly adjoining parietal operculum). Building on fMRI-constrained ERP source analyses of the target P3b (Bledowski et al., 2004), considering that CPP and P3b may reflect the same process (Twomey et al., 2016), and on forward-model-informed regression of scalp topographies onto anatomically defined cortical sources (Mohr et al., 2024), we provide, to our knowledge, the first CPP source estimates derived from subject-specific finite-element volume conduction models with individually digitised electrode positions and fMRI-derived spatial priors. Our source modeling results resonate with and extend previous efforts to pinpoint the generators of the CPP-like family of signals. Prior EEG-fMRI studies and intracranial EEG recordings have indicated that distributed cortical sources give rise to the evidence accumulation signal (e.g., Gherman et al. (2024), Pisauro et al. (2017), Xie et al. (2024), Goueytes et al. (2025) and Stockart et al. (2025)). Our data are generally in line with this distributed-source picture, in that the fit to the CPP topography continued to improve with a large number of added sources, while also singling out certain regions such as the left inferior parietal cortex and right operculum, which appear to contribute most consistently

Beyond the parietal cortex, our localization of an opercular source for the CPP underscores the involvement of the opercular control network in the latter stages of decision formation. This finding dovetails with the evidence that the opercular network shows a spike of activity at decision resolution, especially for uncertain decisions (Pierce et al., 2023). In our study, even though decisions were made on simple target detection, eventually, the opercular regions may still engage to finalize the decision and perhaps initiate performance monitoring. The right opercular focus might indicate a right-lateralized component of this network.

The CPP appears as a symmetric, midline-focused potential on the scalp, whereas the sources identified in our fMRI data were located quite laterally. It is thus not surprising that several sources had to be included together, including regions in both hemispheres, before the predicted, forward-modelled topography attained a symmetry similar to the observed CPP (Fig. 4). It is interesting, however, that the fMRI activations were not themselves perfectly symmetric and balanced, with the left hemisphere predominating. It may be that our multimodal modeling reveals subtle source asymmetries that conventional scalp-level EEG alone would mask. In general, a centrally-distributed scalp ERP can arise from joint projected activity contributed from mirror-image regions of the left and right hemispheres – or in some cases, from a unilateral source whose field spreads broadly (Johnson, 1988; Kičić et al., 2008). We interpret our results as indicating that both hemispheres contribute to the CPP, but the left hemisphere’s contribution was somewhat stronger or more focal such that the source localization procedure picked it up. If, for example, the left parietal cortex had a slightly larger signal than the right, the combined scalp field would still be maximal at the midline (since the voltages from left and right parietal sources summate vectorially and appear centrally). Minor asymmetries in source strength can be masked on the scalp by the smoothing effect of the head’s volume conduction (Abbond et al., 1996). In our analysis, using high-resolution FEM head model and fMRI priors, we were able to detect that the left parietal/right opercular sources best explained the scalp data. Indeed, it is possible that the homologous right parietal/left opercular regions were also active, but did not activate sufficiently to be detected robustly in the initial conjunction analysis.

Another relevant factor is that our participants all responded with their right hand, which would engage the left-hemisphere motor and premotor regions more strongly. Because the decision circuit ultimately has to drive a unimanual motor response, the accumulation process could be more tightly coupled to the contralateral (left) hemisphere sensorimotor areas (Rogge et al., 2022). This would align with findings in monkeys that decision-related build-up can be specific to the hemisphere controlling the action (Cui & Andersen, 2007). Yet, we observed that similar brain regions contributed to scalp-level EEG in mental counting with no motor execution. Another possibility could be the left-hemispheric specialization for certain cognitive processes if our task involved categorical judgment e.g., the left hemisphere often dominates in tasks requiring categorical decision or language-related processing (Dehaene & Cohen, 2007). In sum, a combination of coordinated specialized brain regions could tilt the balance to the left side, even though the overall decision network is bilateral.

Gain- and ablation-based analyses converged on a similar subset of fMRI-informed sources while measuring complementary aspects of model contribution. Whereas gain reflects the fitted weighting of each source projection in the joint reconstruction and can be influenced by covariance among predictors, leave-one-cluster-out ΔMSE quantifies the loss of fit that cannot be recovered by refitting the remaining sources. Their strong rank correspondence (ρ = 0.87; four shared Top-5 clusters) therefore supports the robustness of the leading source subset. Moreover, the first five ranked clusters preferentially recovered the characteristic CPP field while substantial residual error remained at frontal electrodes, even though whole-scalp MSE continued to improve as additional sources were included. This spatial selectivity suggests that the leading parietal–opercular sources are particularly relevant to generation of the CPP rather than simply improving the scalp fit globally. Nevertheless, these results remain model-dependent: the FEM leadfield is derived from structural anatomy independently of the measured CPP and therefore does not itself introduce direct EEG circularity, but the candidate space is constrained by fMRI and source ranking and top-K reconstruction are presently assessed on the same EEG data. We therefore interpret these sources as plausible CPP generators and the top-K analysis as convergent, rather than independent, localization evidence.

### 4.3 Counting without motor action highlights the operculum as the largest CPP contributor

Because the button-press CPP was modelled in a window spanning the response, the right opercular contribution could in principle reflect the manual response rather than decision formation. Indeed, central opercular activation has been reported in a visual oddball task only when targets required a button press (Warbrick et al., 2013). Two further analyses argue against this interpretation. First, we refitted the same forward model to the CPP recorded while participants counted targets covertly, a condition in which the CPP builds without any overt action (O’Connell et al., 2012). The right opercular cortex ranked first by gain and second by unique contribution, and it was the only cluster among the five strongest contributors in both conditions on both measures (Fig. 5). In contrast, the right cerebellar and left postcentral/superior parietal sources, the most plausible carriers of activity related to a right-thumb press, lost their rankings when no response was made.

Second, we refitted the model separately in each participant, using each participant’s own head model. The cluster containing the right opercular cortex had the largest gain during mental counting, and this gain was consistently positive across participants (Fig. S15). Its unique contribution was of similar size with and without a button press, and no cluster contributed more when a response was made. The right opercular contribution is therefore consistent across group-average and individual fits and does not depend on the manual effector. This extends to the source level the effector independence previously shown for the CPP itself (O’Connell et al., 2012; Twomey et al., 2016).

### 4.4 Methodological Considerations and Limitations of FEM Source Modeling

Our fMRI-informed FEM source modeling provides a powerful, anatomically grounded framework for linking distributed decision-related fMRI activations to the scalp CPP, but several caveats apply. EEG source localization is inherently ill-posed, and although subject-specific FEM head models combined with fMRI priors and distributed inverse algorithms improve spatial plausibility, the resulting solutions remain estimates. Accuracy depends on the fidelity of tissue segmentation and conductivity values, particularly in the skull, and orientation constraints favor cortical over deep sources (Montes-Restrepo et al., 2014). Reliance on fMRI priors increases specificity but risks overlooking EEG-relevant activity outside BOLD-active regions, while different inverse approaches (e.g. minimum-norm, beamformers, dipole fits) can yield systematically different solutions (Abreu et al., 2021). Still, individualized FEM models consistently outperform template or simplified geometries, and our results align with known physiology, underscoring the promise of this multimodal approach. Future work using intracranial EEG (e.g., Gherman et al. (2024)) or causal perturbations such as transcranial magnetic stimulation (TMS) (Philiastides et al., 2011) will be critical to validate whether the identified parietal–opercular sources are indeed the generators of the human abstract decision signal.

Our approach can only identify generators that both appeared in the fMRI conjunction and project detectably to the scalp. Regions equally crucial to the decision process may contribute little or no net dipolar current at the scalp — because of closed-field geometry, cancellation across sulcal banks, or depth — and would be invisible here regardless of their functional importance; the gains we report redistribute the observed scalp variance among the modelled sources and do not rank regions by their causal role. Additionally, our task was simple: a single supramodal change detection with a 94.5% hit rate and no manipulation of difficulty, conflict or response mapping. Task complexity is an important dimension for future work to explore. Moreover, our source space was derived from an exploratory, uncorrected conjunction map and is therefore restricted to regions that reached a liberal BOLD criterion. Gains should therefore be read as the best explanation of the observed topography *within this restricted source space*, not as estimates of regional contribution to the CPP.

In summary, combining EEG, fMRI, and individualized head modeling provides a coherent picture of abstract perceptual decision-making: a bilateral parietal-opercular network that accumulates evidence for cognitive control. This multimodal approach sets the foundation for future advances in neural source modeling and causal validation for deeper insight into how the brain transforms sensation into decision.

## 5 Declaration of the use of AI

### Research

AI played no role in study design, data collection, or choice of analyses. ChatGPT (GPT-5, OpenAI) and Claude (Sonnet 5.5 and Opus 5.5, Anthropic) were used as coding assistants to draft and debug parts of the MATLAB code. All AI-assisted code was verified and tested by the authors before use.

### Writing

The same tools were used to check spelling and grammar and to help verify reference details. All citations were checked by the authors against the original sources. No AI was used to generate figures.

## 6 Data and Code Availability

All analysis code used in this study will be made available upon publication. This includes the MATLAB scripts for fMRI analysis, reaction-time analysis, EEG preprocessing, individualised finite-element leadfield computation, ERP/CPP extraction and fMRI-informed forward modelling of the CPP topography. The analyses used MATLAB, SPM12, EEGLAB and FieldTrip, which are freely available from their developers. The raw and processed EEG, fMRI and behavioural data are available from the corresponding authors upon reasonable request, subject to approval by the Trinity College Dublin School of Psychology Ethics Committee and a data-sharing agreement.

## 7 Author Contributions

N.S., S.K., and R.O. designed the experiment. N.S., K.K. acquired the data. A.A., K.W.-L., S.K., and R.O. designed analyses. A.A. performed the analysis, modelling and statistical analyses. A.A., K.W.-L., S.K., and R.O. validated the data and analyses. A.A. wrote the initial draft. K.W.-L., S.K., and R.O. supervised the research.

## 10 Acknowledgements

We are grateful for access to the Tier 2 High-Performance Computing resources provided by the Northern Ireland High-Performance Computing (NI-HPC) facility funded by the UK Engineering and Physical Sciences Research Council (EPSRC), Grant No. EP/T022175/1.

