## Supplementary Materials for "Mapping brain networks underlying abstract perceptual decision formation and its EEG signatures"

The supplementary tables and figures provide additional details on EEG–fMRI source modeling, condition-specific analyses, and validation of our modeling approach.

- **Table S1** summarizes the full set of local maxima identified in the conjunction analysis, reporting cluster sizes, uncorrected cluster-level p-values, and corresponding peak-level T, Z, and uncorrected p-values. This table provides the comprehensive statistical map underlying the subset of peaks used for source analysis.
- **Table S2** lists the peak coordinates from the fMRI conjunction analysis across mental counting and button-press sessions. From clusters exceeding 10 voxels, we selected 24 local maxima, primarily from the first and second clusters, to define the priors for EEG source modeling.
- **Figure S1** shows trial-wise and grand-average centroparietal positivity (CPP) patterns in the separate EEG dataset (N = 31), illustrating stronger CPP build-up in auditory-up and visual-up conditions compared to their respective down conditions.
- **Figure S2** illustrates that decision-related brain sources contribute to the scalp decision signal, showing grand-average CPP waveforms and topographic maps separately for fast, medium, and slow RT bins across stimulation conditions.
- **Figures S3–S6** present **single-source contribution analyses** for each stimulation condition (AU, AD, VU, VD), highlighting the dominant fMRI-informed sources projecting to centroparietal electrodes.
- **Figures S7–10** extend this to **pairwise source contribution analyses**, testing the combined explanatory power of bilateral cortical sources across conditions.

- **Figure S11** depicts **projected scalp topographies** for each individual fMRI-derived source, illustrating how each contributes to the observed EEG distribution at the scalp.
- **Figure S12** shows the anatomical locations of the top-ranked fMRI-derived dipole sources rendered on a three-dimensional brain mesh, providing a spatial overview of the cortical regions most strongly contributing to the centroparietal EEG signal across conditions.

- **Table S3** ranks the fMRI-derived source clusters by their mean absolute fitted gain across seven response-locked timepoints, reporting their constituent sources, MNI coordinates, and normalized gain values. This table provides a quantitative assessment of the relative contributions of individual source clusters to the reconstructed scalp EEG topography.

- **Figure S13** presents the ablation-based ranking of source clusters, evaluating their unique contributions to model fit by measuring the increase in mean squared error (ΔMSE) following the removal of each cluster. The figure illustrates the relative importance of individual clusters in explaining the observed scalp EEG activity across the response-locked time window.

- **Table S4** summarizes the ablation-based ranking of fMRI-derived source clusters, reporting their constituent sources, MNI coordinates, and corresponding ΔMSE values. This table complements the gain-based ranking in Table S3 by quantifying the unique contribution of each cluster to the accuracy of the reconstructed scalp EEG topography.

- **Figure S14** shows the centroparietal positivity (CPP) recorded during mental counting without a manual response. It gives the target-locked CPP waveform, which peaked at 869 ms, and the evolution of its scalp topography; this peak defined the analysis window for the motor-free forward-model fit.
- **Figure S15** presents the participant-level forward-model fits, in which each participant's own head model and topographies were used with 10 well-conditioned source clusters. It shows the per-participant gain and ablation for both conditions, with permutation-based family-wise-error-corrected inference; the cluster containing the right opercular cortex was the strongest contributor during mental counting.
- **Table S5** lists the composition of the 10 source clusters used in the participant-level analysis, with the anatomical labels and MNI coordinates of their member sources.
- **Figure S16** compares observed and model-predicted grand-mean topographies for button press and mental counting, using the 24 raw sources and the 10 clusters. It shows that clustering preserved the main features of the CPP topography, with little loss of spatial correspondence.

Together, these tables and figures provide converging evidence for the supramodal and spatially consistent contributions of parietal and precentral sources to the CPP across sensory modalities.

**Table S1.** Peak coordinates from the fMRI conjunction analysis across mental counting and manual button-press tasks. The table lists all local maxima (uncorrected peak-level p < 0.05) along with cluster size, cluster-level p-values, and corresponding T and Z statistics. Reported peaks represent regions consistently recruited during perceptual decision-making irrespective of modality (auditory/visual) or motor requirement (counting/button press).

| MNI coordinates (x, y, z) | Peak-level p (unc.) | Peak T | Peak $Z_E$ | Cluster size (voxels) | Cluster-level p (unc.) |
|---|---|---|---|---|---|
| (60, -16, 23) | 0 | 3.98 | 3.9 | 926 | 0 |
| (45, -37, 29) | 0 | 3.79 | 3.72 | | |
| (51, -28, 56) | 0 | 3.6 | 3.54 | | |
| (-57, 5, 5) | 0 | 3.74 | 3.68 | 176 | 0.064 |
| (-45, 8, -1) | 0.002 | 2.86 | 2.83 | | |
| (-60, 5, 26) | 0.003 | 2.81 | 2.78 | | |
| (-63, -19, 26) | 0 | 3.42 | 3.38 | 411 | 0.008 |
| (-51, -13, 53) | 0.001 | 3.28 | 3.24 | | |
| (-54, -13, 41) | 0.001 | 3.08 | 3.05 | | |
| (-48, -40, 56) | 0.001 | 3.19 | 3.16 | 44 | 0.335 |
| (54, 5, 41) | 0.002 | 2.97 | 2.94 | 39 | 0.365 |
| (24, -58, -19) | 0.006 | 2.55 | 2.53 | 19 | 0.587 |
| (-39, -34, 23) | 0.035 | 1.81 | 1.81 | 1 | 0.916 |
| (21, -46, 59) | 0.039 | 1.76 | 1.76 | 1 | 0.916 |
| (39, -34, 35) | 0.04 | 1.75 | 1.75 | 1 | 0.916 |
| (24, -40, 50) | 0.041 | 1.74 | 1.74 | 2 | 0.87 |
| (-48, -10, 26) | 0.043 | 1.72 | 1.72 | 1 | 0.916 |
| (27, -37, 47) | 0.043 | 1.71 | 1.71 | 1 | 0.916 |
| (54, -7, 47) | 0.047 | 1.67 | 1.68 | 1 | 0.916 |
| (-21, -43, 56) | 0.047 | 1.67 | 1.67 | 1 | 0.916 |
| (33, -37, 35) | 0.047 | 1.67 | 1.67 | 2 | 0.87 |

**Table S2. Selected peak coordinates from the fMRI conjunction analysis.** From clusters larger than 10 voxels in Table S1, we retained 24 local maxima, predominantly originating from the two largest clusters, to serve as priors for the EEG source analysis.

| Brain region | MNI coordinates (x, y, z) | Cluster size (voxels) | Cluster-level p (unc.) | Peak T | Pea k Z | Peak-level p (unc.) |
|---|---|---|---|---|---|---|
| rPostCentral Gyrus | (60, -16, 23) | 926 | 0 | 3.98 | 3.9 | 0 |
| rPostCentral Gyrus (rSPL) | (45, -37, 59) | | | | | |
| rPostCentral Gyrus | (51, -28, 56) | | | | | |
| rSupramarginal Gyrus (rIPL) | (63, -37, 26) | | | | | |
| rPostCentral Gyrus | (48, -31, 59) | | | | | |
| rSupramarginal Gyrus (rIPL) | (57, -25, 50) | | | | | |
| rSupramarginal Gyrus (rIPL) | (66, -25, 32) | | | | | |
| rPrecentral Gyrus | (63, 8, 8) | | | | | |
| rPostCentral Gyrus | (63, -16, 41) | | | | | |
| rOpercular Cor. | (48, 2, 5) | | | | | |
| lPostCentral Gyrus | (-63, -19, 26) | 411 | 0.008 | 3.42 | 3.38 | 0.001 |
| lPrecentral Gyrus | (-51, -13, 53) | | | | | |
| lPostCentral Gyrus | (-63, -13, 23) | | | | | |
| lPostCentral Gyrus | (-54, -13, 41) | | | | | |
| lSupramarginal Gyrus (lIPL) | (-63, -31, 23) | | | | | |
| lLateral Occ. | (-54, -67, 5) | | | | | |
| lSupramarginal Gyrus (lIPL) | (-48, -43, 26) | | | | | |
| lSupramarginal Gyrus (lIPL) | (-54, -46, 29) | | | | | |
| lSupramarginal Gyrus (lIPL) | (-54, -46, 35) | | | | | |
| lPostCentral Gyrus | (-45, -37, 32) | | | | | |
| lPrecentral Gyrus | (-57, 5, 5) | 176 | 0.064 | 3.74 | 3.68 | 0 |
| lPostCentral Gyrus (lSPL) | (-48, -40, 56) | 44 | 0.335 | 3.19 | 3.16 | 0.001 |
| rPrecentral Gyrus (rMFG) | (54, 5, 41) | 39 | 0.365 | 2.97 | 2.94 | 0.002 |
| rCerebellum | (24, -58, -19) | 15 | 0.587 | 2.55 | 2.53 | 0.006 |

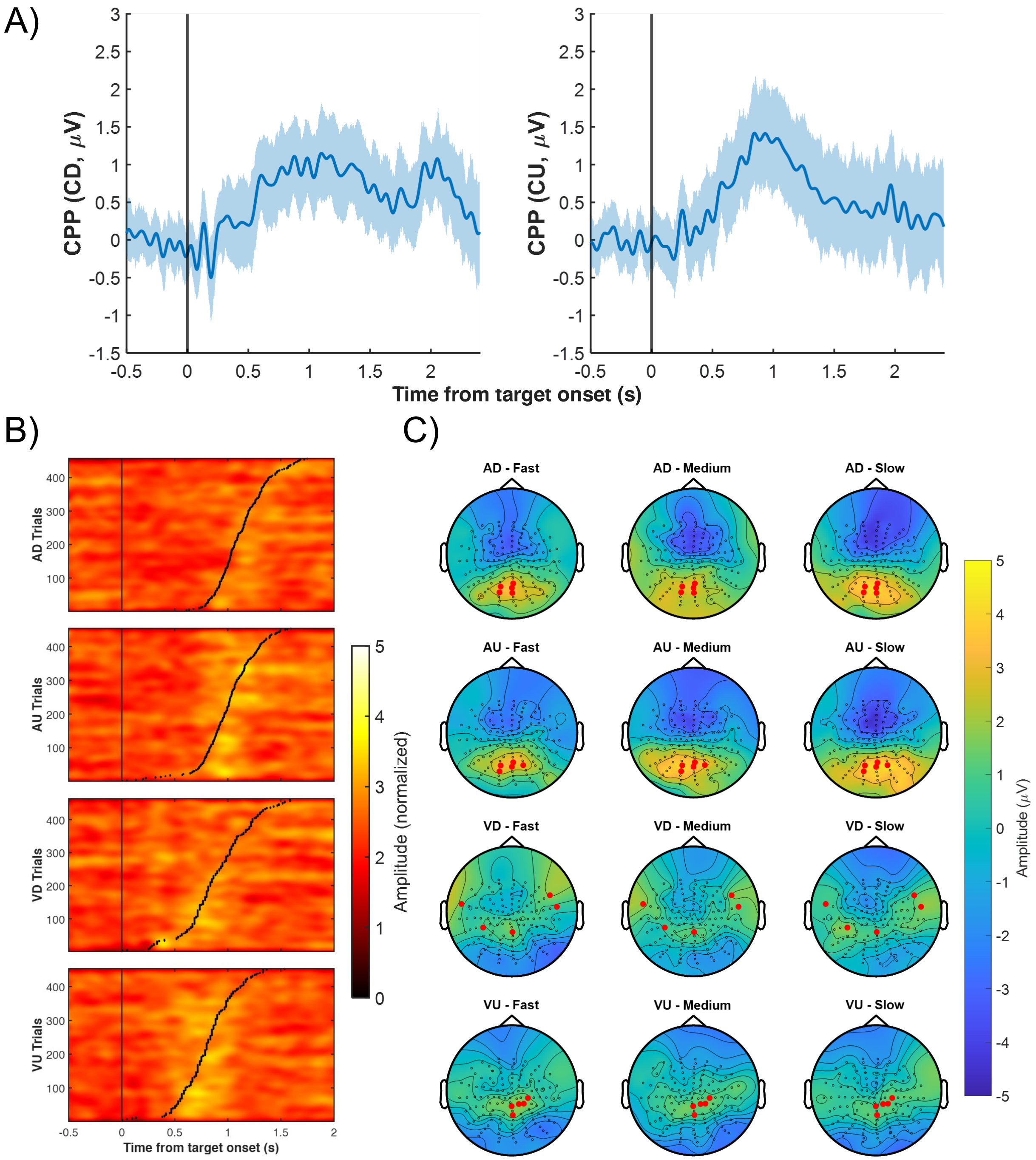


**Figure S1. CPP across conditions in the separate EEG dataset (N = 31).** A) Averaged centroparietal positivity (CPP) during the mental counting session, derived from predefined centroparietal electrodes, shows a sustained positivity that evolves gradually and peaks in the counting visual up (CU) and counting visual down (CD) conditions. B) trial-by-trial stimulus-locked CPP activity, pooled across participants, baseline-corrected, normalized to [0–5], and sorted within condition by each trial's reaction time (RT), marked by black dots. Higher-amplitude CPP ramps are evident around RTs for auditory-up (AU) and visual-up (VU) conditions, whereas auditory-down (AD) and visual-down (VD) show noticeably weaker CPP expression over the same window. C) Topographic maps showing amplitude in the window X-Yms relative to response in each of three equal RT bins, grand averaged across all participants. Although apparently weaker in visual conditions, possibly due to overlapping occipital negative activity, all conditions include a centroparietal focus.

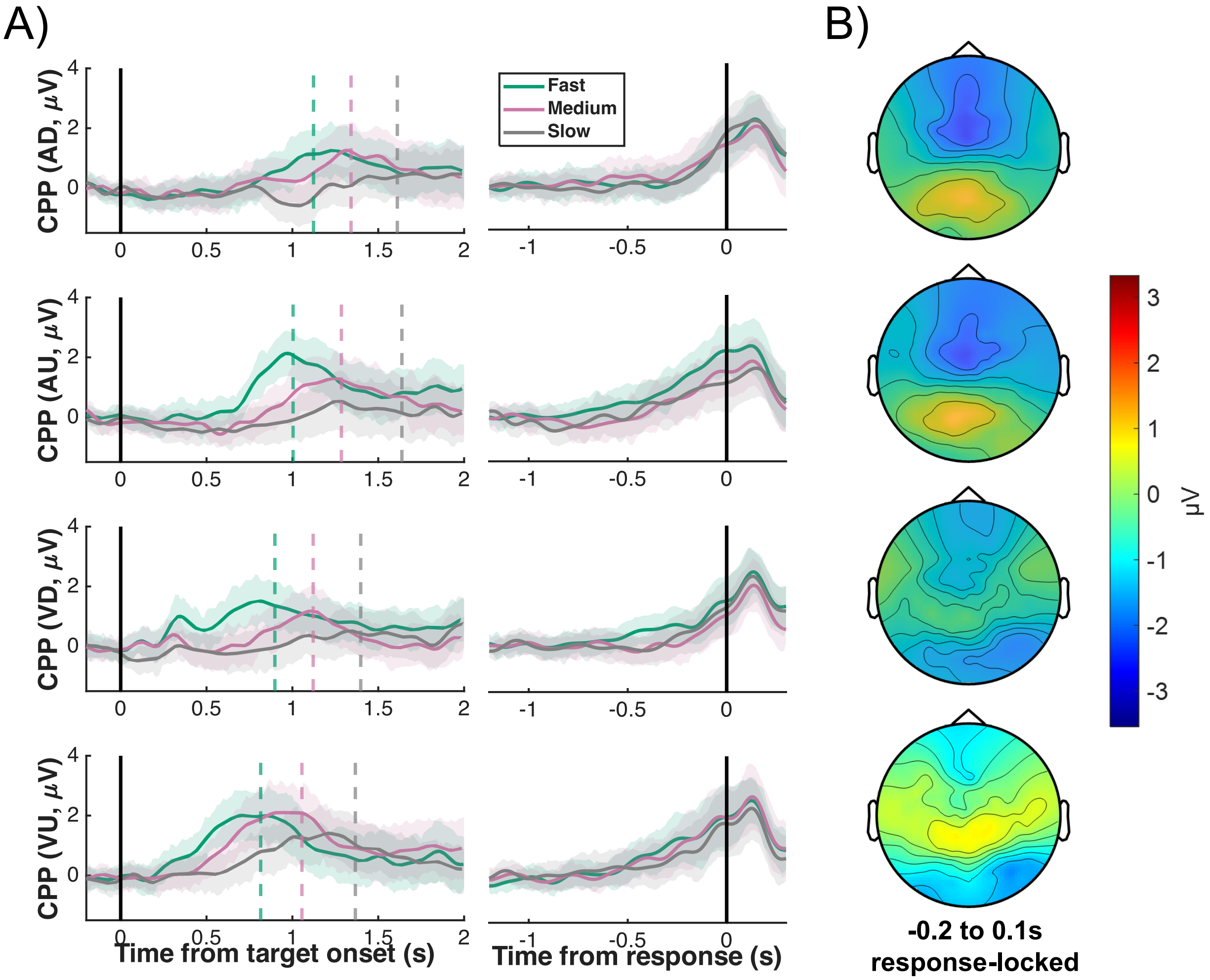


**Figure S2. Decision-related brain sources contribute to the scalp decision signal.** A) Grand average of CPP activity over participants in different conditions during button press sessions for trials with fast, medium, and slow reactions. B) Corresponding topographic maps of response-locked EEG activity averaged from −200 ms to 100 ms from button press, separately for each stimulation. Abbreviations as in Figure S1.

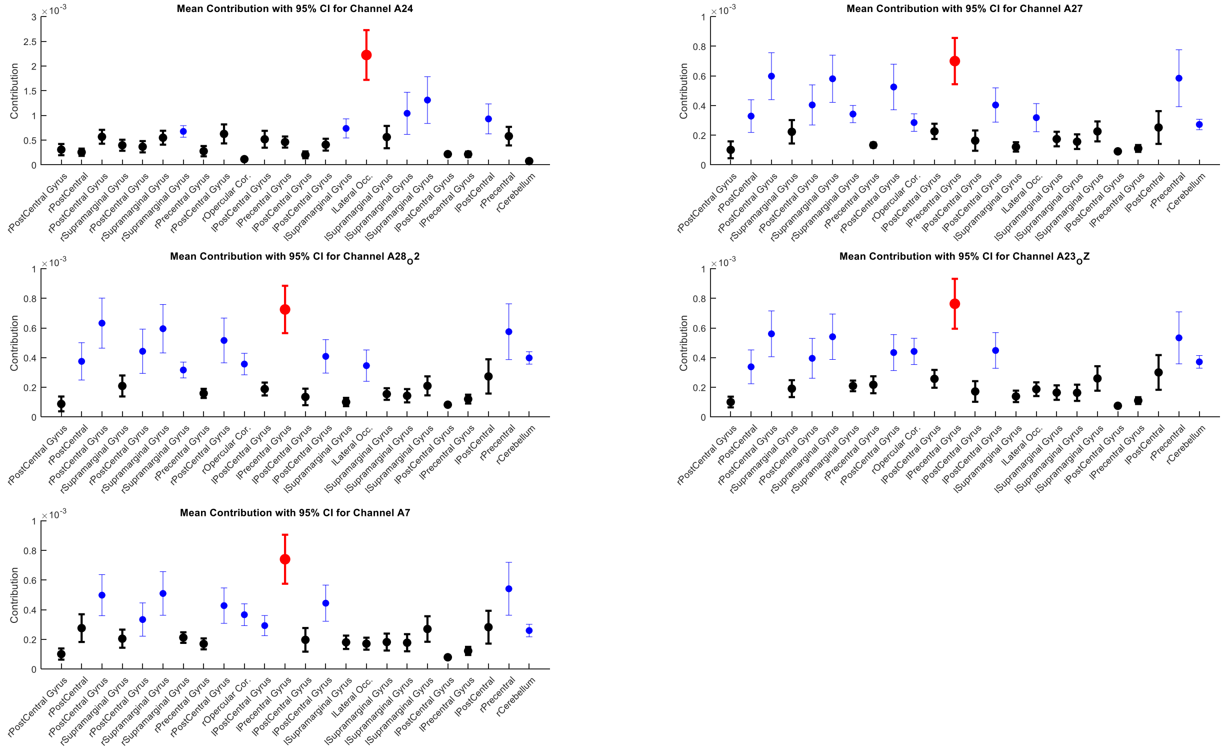


**Figure S3. Single-source contribution analysis for the auditory up condition.** In this condition, the left precentral gyrus consistently emerged as the dominant contributor across electrodes. Each subplot shows the average contribution of individual fMRI-informed sources to one of the five centroparietal electrodes with highest amplitude at −50 ms (region-of-interest (ROI)). Bars represent the mean contribution across participants; error bars indicate 95% confidence intervals. The source with the highest average contribution per electrode is marked in red, while statistically significant differences (relative to the top source) based on Bonferroni-corrected post hoc tests are marked in black. All other sources are shown in blue.

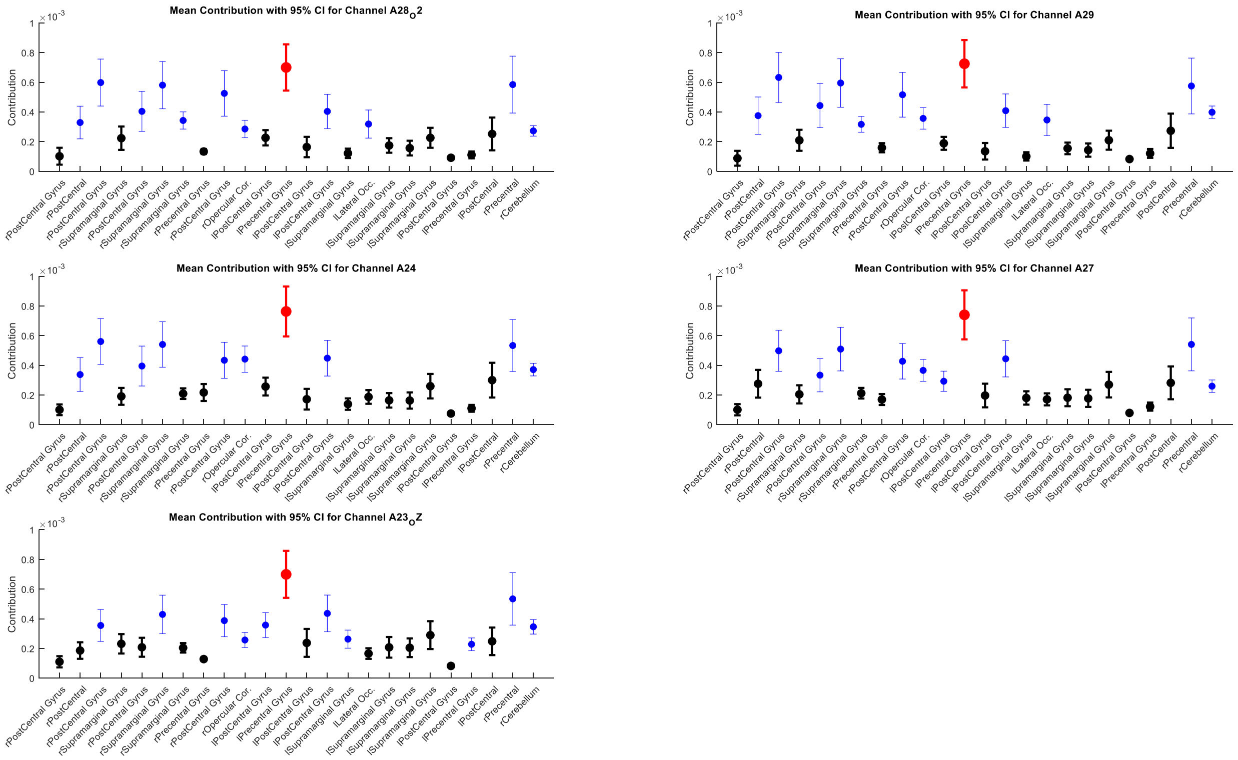


**Figure S4. Single-source contribution analysis for the auditory down condition.** Dominant sources across ROI electrodes largely localized to the left precentral gyrus, mirroring the auditory up pattern. Full explanation of bar color coding and statistical conventions is provided in the Figure S3 caption.

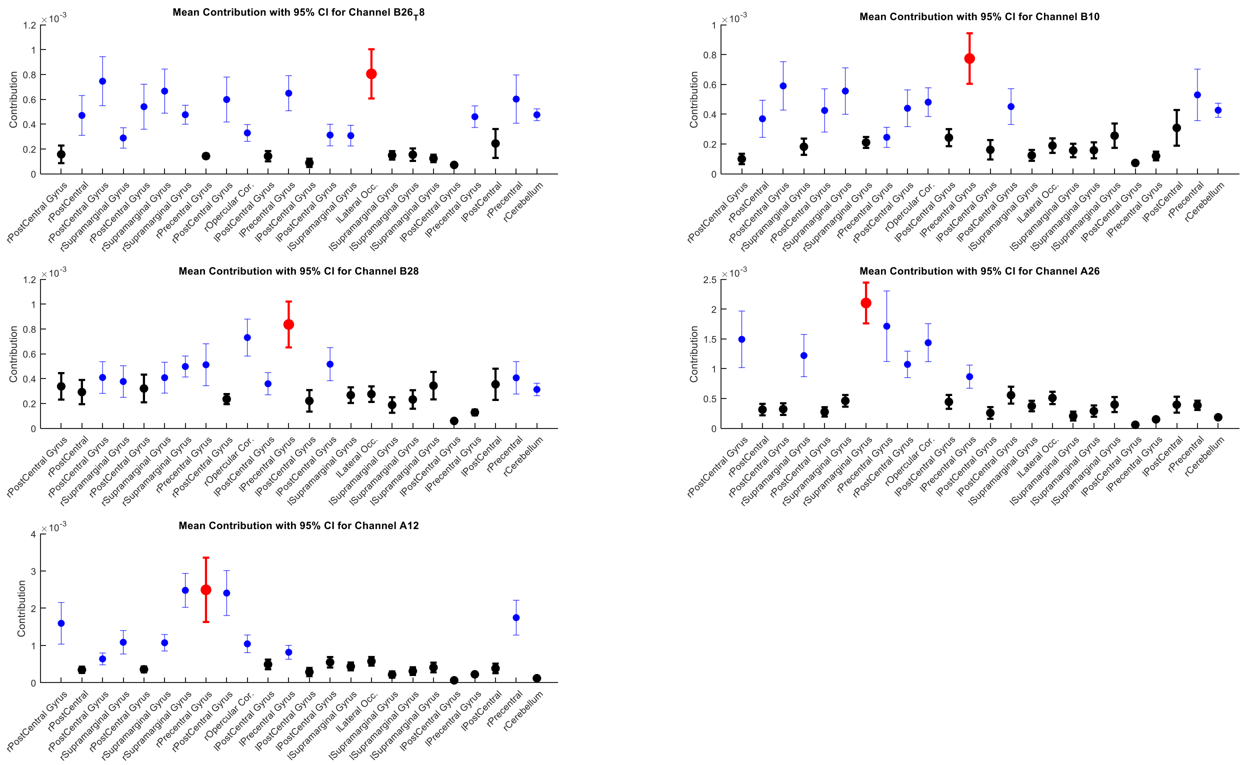


**Figure S5. Single-source contribution analysis for the visual up condition.** Contributions were more bilaterally distributed, with both left and right precentral gyri emerging as high-contributing regions across centroparietal electrodes. Full explanation of bar color coding and statistical conventions is provided in the Figure S3 caption.

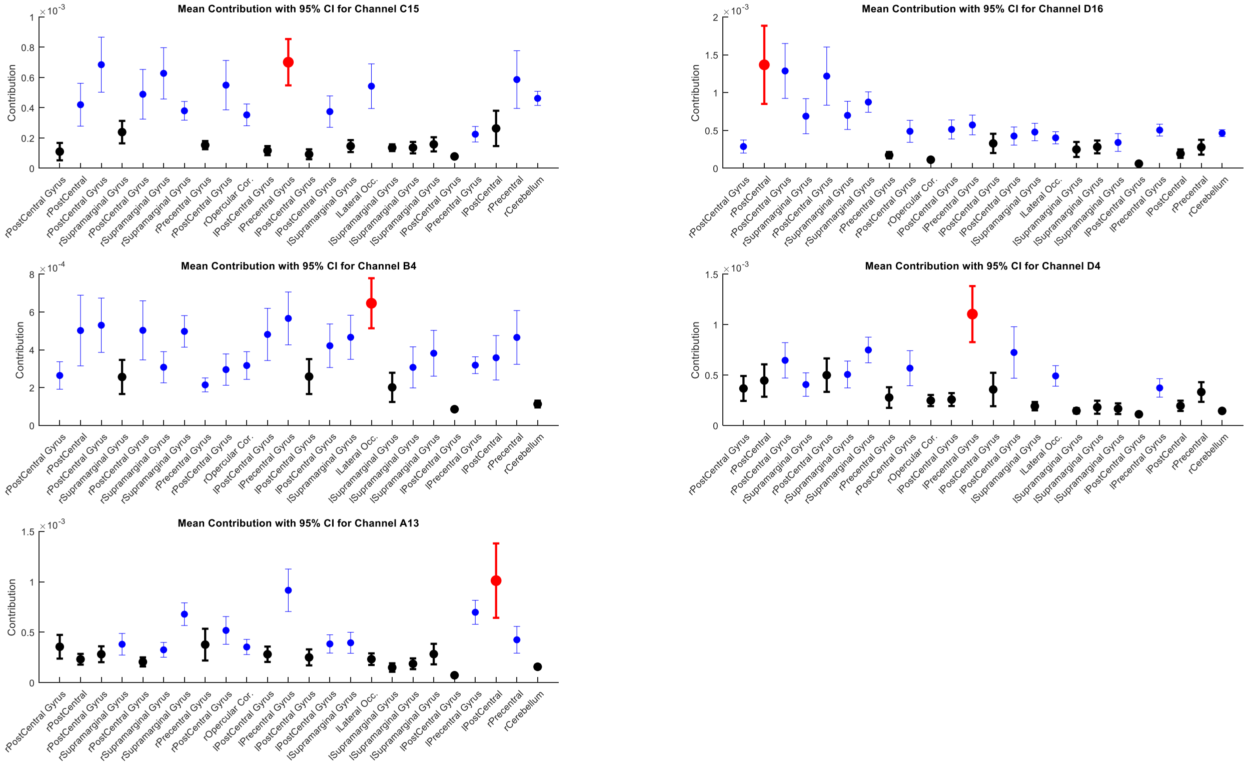


**Figure S6. Single-source contribution analysis for the visual down condition.** Unlike other conditions, the visual down task yielded a more heterogeneous distribution of contributions without a consistent dominant source, suggesting broader or less centralized cortical involvement. Full explanation of bar color coding and statistical conventions is provided in the Figure S3 caption.

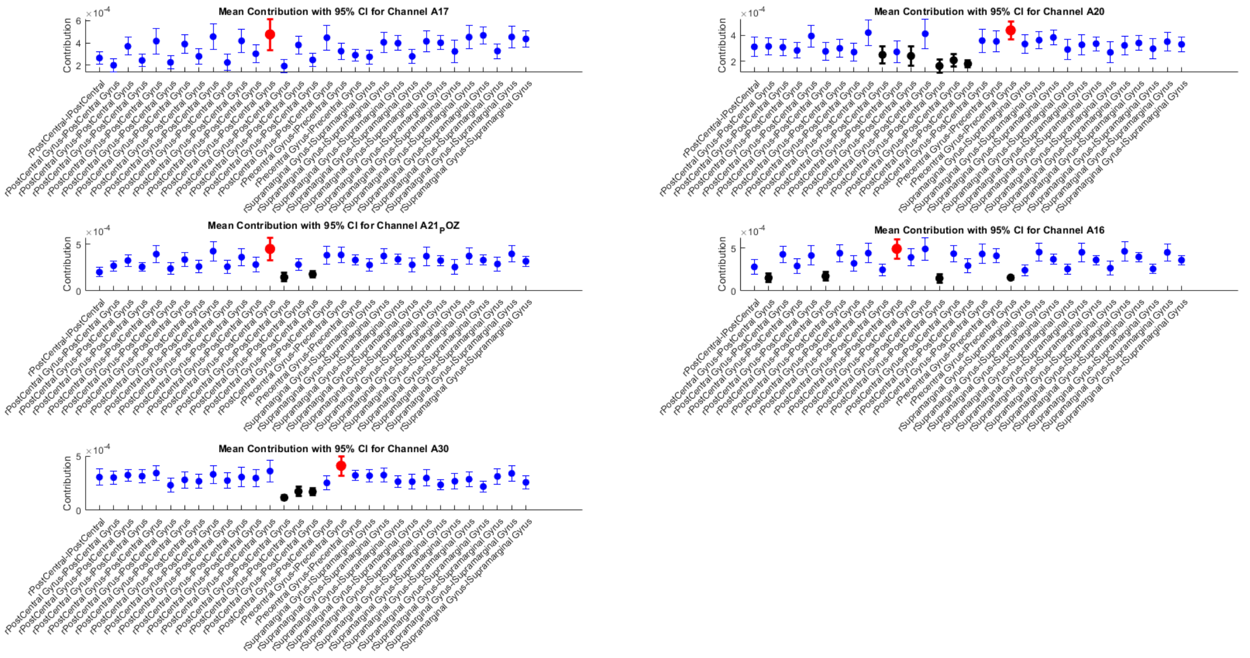


**Figure S7. Pairwise source contribution analysis for the auditory up condition.** Left–right postcentral gyrus consistently emerged as the top-contributing pair across electrodes. Each subplot corresponds to one of the five ROI electrodes with the highest amplitude at −50 ms. Error bars represent 95% confidence intervals across participants. The source pair with the highest mean contribution is marked in red; statistically significant differences from this top pair (Bonferroni-corrected) are marked in black; all other pairs are in blue.

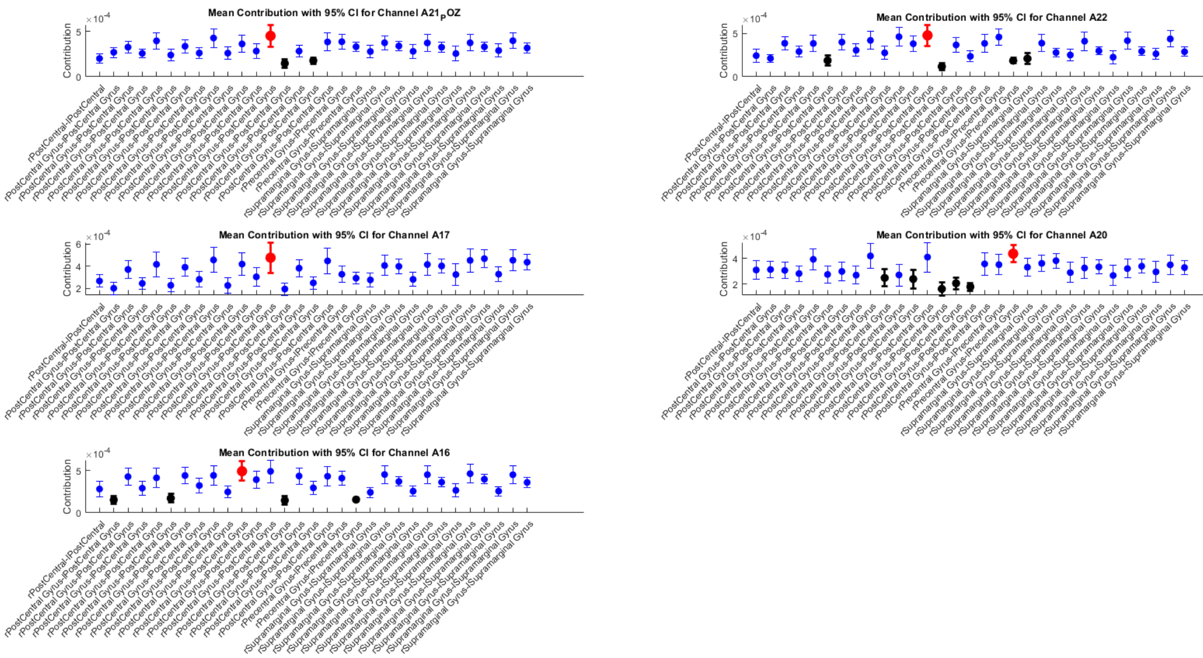


**Figure S8. Pairwise source contribution analysis for the auditory down condition.** Statistical comparisons of all anatomically defined source pairs across ROI electrodes reveal the dominant role of bilateral postcentral gyrus contributions. Full explanation of bar color coding and statistical conventions is provided in the Figure S7 caption.

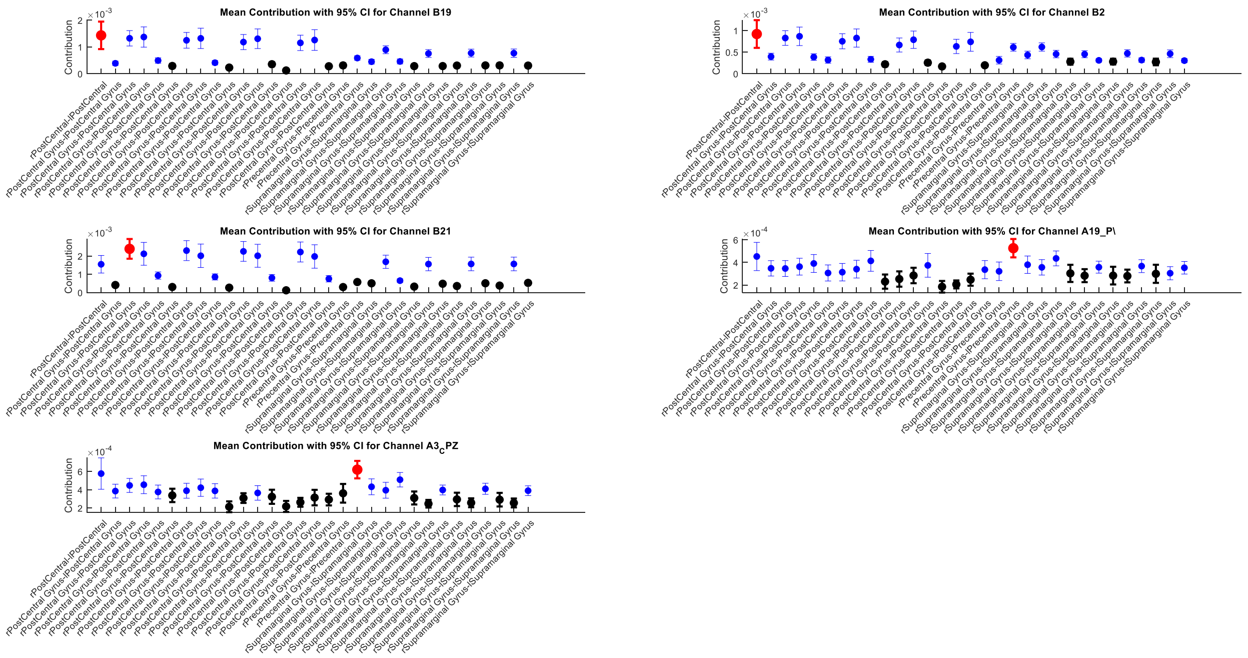


**Figure S9. Pairwise source contribution analysis for the visual up condition.** Bilateral postcentral sources again contributed most strongly, though peak pairs varied slightly across electrodes. Full explanation of bar color coding and statistical conventions is provided in the Figure S7 caption.

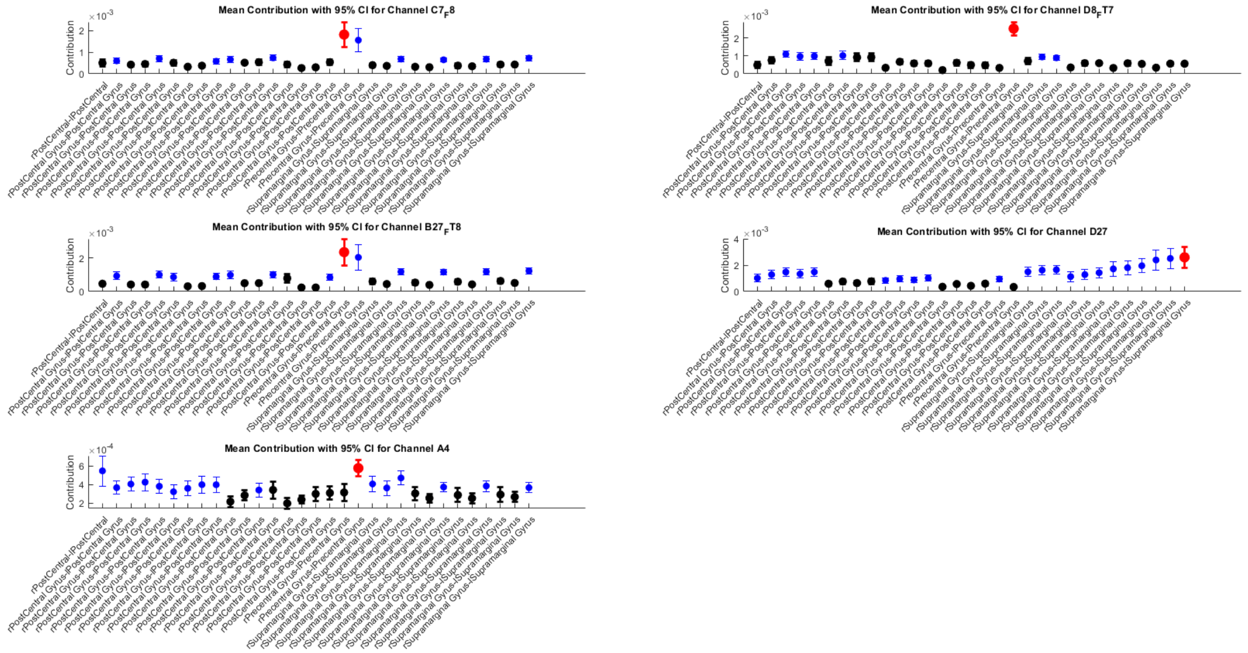


**Figure S10. Pairwise source contribution analysis for the visual down condition.** The visual down task revealed dominant contributions from bilateral precentral gyrus pairs. Full explanation of bar color coding and statistical conventions is provided in the Figure S7 caption.

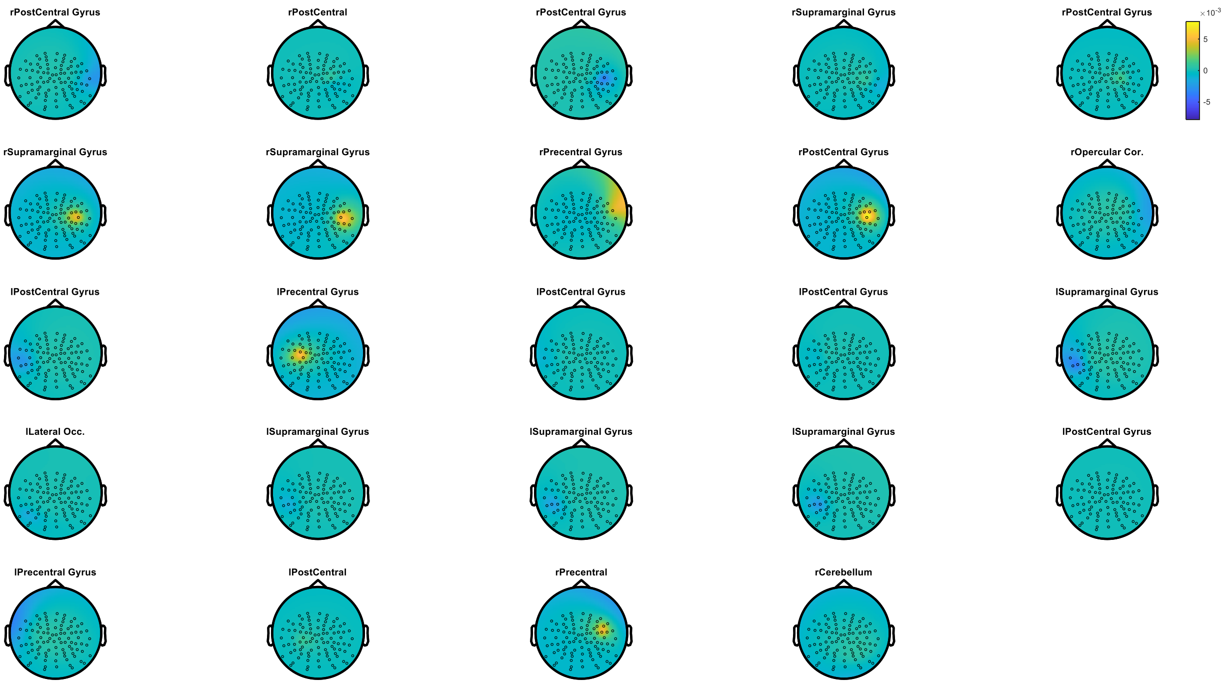


**Figure S11. Projected scalp activity (predictor) from each fMRI-derived source.** Each panel displays the modeled topography resulting from the projection of a single dipole source onto the scalp electrode array, using averaged participant leadfields and canonical dipole orientations.

**Table S3. Clusters ranked by gain magnitude, with constituent source coordinates.** Clusters are ranked (Rank 1 = highest) by mean absolute gain across the seven response-locked timepoints analyzed (−100 to +50 ms, 25 ms steps; see Methods). "Cluster Members" lists the constituent fMRI-derived source(s) merged into each cluster during correlation-based dimensionality reduction (see Methods; full peak-level details in Table S2), with corresponding MNI coordinates given under "Members MNI Coordinates," in the same order. Where a cluster combines multiple constituent sources (e.g. Rank 1, 6, 9, 14, 16), its gain reflects the fitted contribution of the merged (averaged) predictor topography, not the sum of its members' individual contributions.

| Rank | Cluster Members | Members MNI Coordinates | Gain ± SD | Normalised Gain ± SD |
|---|---|---|---|---|
| 1 | lPostCentral Gyrus, lPostCentral Gyrus | (-54, -13, 41); (-45, -37, 32) | 1.18 ± 0.27 | 0.78 ± 0.18 |
| 2 | rOpercular Cor. | (48, 2, 5) | 1.01 ± 0.20 | 0.66 ± 0.14 |
| 3 | rCerebellum | (24, -58, -19) | 0.71 ± 0.12 | 0.46 ± 0.08 |
| 4 | lPostCentral | (-48, -40, 56) | 0.58 ± 0.17 | 0.38 ± 0.11 |
| 5 | rPostCentral Gyrus | (60, -16, 23) | 0.43 ± 0.08 | 0.28 ± 0.05 |
| 6 | lPostCentral Gyrus, lPostCentral Gyrus, lSupramarginal Gyrus | (-63, -19, 26); (-63, -13, 23); (-63, -31, 23) | 0.35 ± 0.06 | 0.23 ± 0.04 |
| 7 | lLateral Occ. | (-54, -67, 5) | 0.29 ± 0.08 | 0.19 ± 0.05 |
| 8 | rSupramarginal Gyrus | (63, -37, 26) | 0.29 ± 0.04 | 0.19 ± 0.03 |
| 9 | lSupramarginal Gyrus, lSupramarginal Gyrus, lSupramarginal Gyrus | (-48, -43, 26); (-54, -46, 29); (-54, -46, 35) | 0.22 ± 0.03 | 0.14 ± 0.02 |
| 10 | rPrecentral Gyrus | (63, 8, 8) | 0.21 ± 0.09 | 0.14 ± 0.06 |
| 11 | rPostCentral Gyrus | (48, -31, 59) | 0.20 ± 0.05 | 0.13 ± 0.03 |
| 12 | rPostCentral | (45, -37, 59) | 0.15 ± 0.03 | 0.09 ± 0.02 |
| 13 | rPrecentral | (54, 5, 41) | 0.14 ± 0.03 | 0.08 ± 0.02 |
| 14 | rPostCentral Gyrus, rSupramarginal Gyrus | (51, -28, 56); (66, -25, 32) | 0.13 ± 0.02 | 0.08 ± 0.01 |
| 15 | lPrecentral Gyrus | (-51, -13, 53) | 0.13 ± 0.05 | 0.08 ± 0.03 |
| 16 | rSupramarginal Gyrus, rPostCentral Gyrus | (57, -25, 50); (63, -16, 41) | 0.08 ± 0.03 | 0.05 ± 0.02 |
| 17 | lPrecentral Gyrus | (-57, 5, 5) | 0.03 ± 0.04 | 0.01 ± 0.02 |

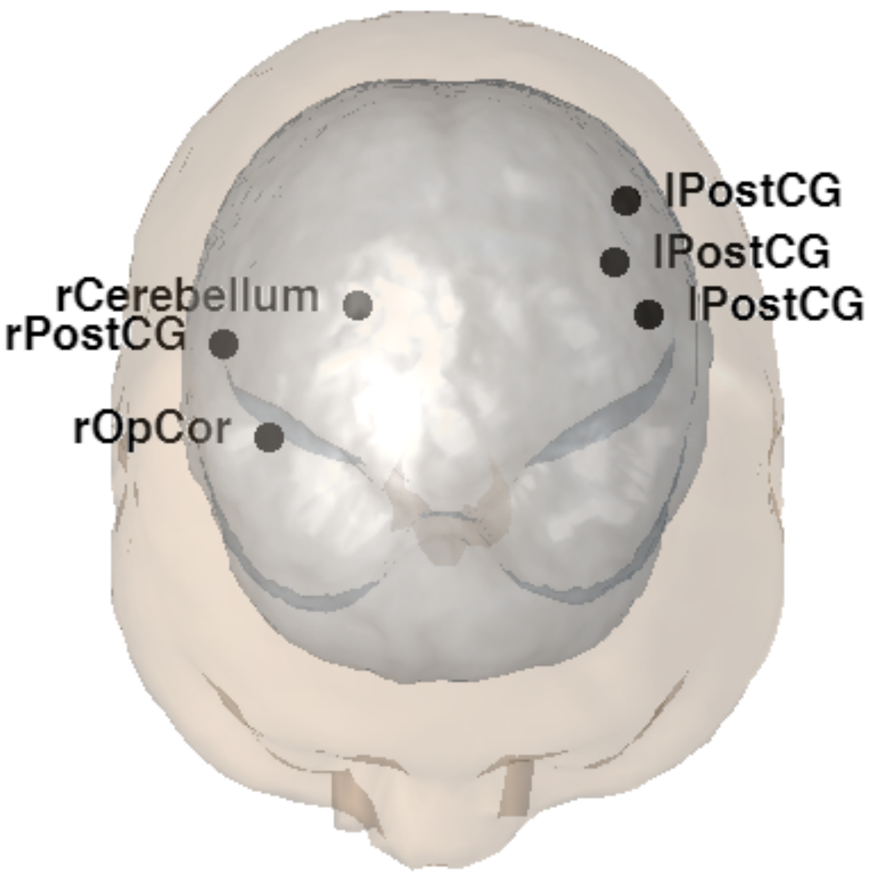


**Figure S12. Anatomical localisation of top-ranked fMRI-informed dipole sources on a three-dimensional brain mesh.** Black filled circles indicate the MNI coordinates of the fMRI-derived dipole sources that contributed most strongly to the centroparietal EEG signal across conditions. The underlying meshes show the cortical surface (blue-grey, semi-transparent) and the scalp surface (beige, semi-transparent). The brain is displayed from a posterior-superior viewpoint (azimuth 180°, elevation 55°). Source abbreviations: lPostCG = Left Postcentral Gyrus; rPostCG = Right Postcentral Gyrus; rOpCor = Right Opercular Cortex; rCerebellum = Right Cerebellum.

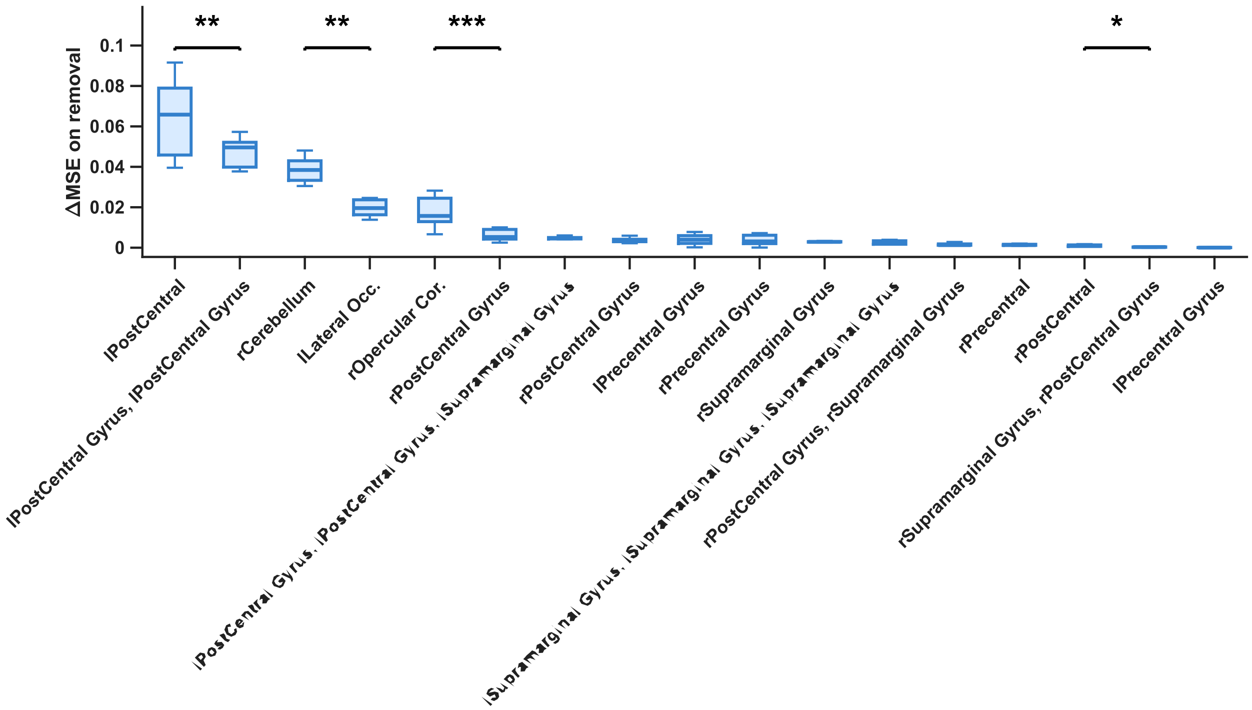


**Figure S13. Clusters ranked by ablation importance.** For each cluster, the model was refit with that cluster excluded, and ΔMSE (= Mean Squared Error (MSE) without the cluster − MSE of the full 17-cluster model) was computed separately at each of the seven response-locked timepoints analyzed (−100 to +50 ms, 25 ms steps; see Methods). Boxplots show the resulting distribution of ΔMSE across these seven timepoints for each cluster, ranked in descending order of mean ΔMSE (leftmost = greatest increase in error upon removal, i.e. greatest unique contribution to model fit). Significance brackets compare adjacent-ranked clusters (Lilliefors test for normality of the paired differences, followed by a paired t-test or Wilcoxon signed-rank test as appropriate; *p < 0.05, **p < 0.01, ***p < 0.001).

**Table S4. Clusters ranked by ablation importance, with constituent source coordinates.** Clusters are ranked (Rank 1 = greatest change in mean square error (ΔMSE) upon removal) by mean ΔMSE across the seven response-locked timepoints analyzed (−100 to +50 ms, 25 ms steps; see Methods). "Cluster Members" lists the constituent fMRI-derived source(s) merged into each cluster during correlation-based dimensionality reduction (see Methods; full peak-level details in Table S2), with corresponding MNI coordinates given under "Members MNI Coordinates," in the same order (identical cluster composition to Table S3, reordered by ablation rank). "ΔMSE ± SD" reports each cluster's mean ΔMSE ± standard deviation across the seven timepoints. "Normalised ΔMSE ± SD" reports the same values after min-max normalization to the [0, 1] range, computed jointly across all clusters and timepoints.

| Rank | Cluster Members | Members MNI Coordinates | ΔMSE ± SD | Normalized ΔMSE ± SD |
|---|---|---|---|---|
| 1 | lPostCentral | (-48, -40, 56) | 0.06 ± 0.02 | 0.70 ± 0.21 |
| 2 | lPostCentral Gyrus,<br>lPostCentral Gyrus | (-54, -13, 41);<br>(-45, -37, 32) | 0.05 ± 0.01 | 0.51 ± 0.08 |
| 3 | rCerebellum | (24, -58, -19) | 0.04 ± 0.01 | 0.42 ± 0.07 |
| 4 | lLateral Occ. | (-54, -67, 5) | 0.02 ± 0.00 | 0.22 ± 0.05 |
| 5 | rOpercular Cor. | (48, 2, 5) | 0.02 ± 0.01 | 0.19 ± 0.08 |
| 6 | rPostCentral Gyrus | (60, -16, 23) | 0.01 ± 0.00 | 0.07 ± 0.03 |
| 7 | lPostCentral Gyrus,<br>lPostCentral Gyrus,<br>lSupramarginal Gyrus | (-63, -19, 26);<br>(-63, -13, 23);<br>(-63, -31, 23) | 0.00 ± 0.00 | 0.05 ± 0.01 |
| 8 | rPostCentral Gyrus | (48, -31, 59) | 0.00 ± 0.00 | 0.04 ± 0.02 |
| 9 | lPrecentral Gyrus | (-51, -13, 53) | 0.00 ± 0.00 | 0.04 ± 0.03 |
| 10 | rPrecentral Gyrus | (63, 8, 8) | 0.00 ± 0.00 | 0.04 ± 0.03 |
| 11 | rSupramarginal Gyrus | (63, -37, 26) | 0.00 ± 0.00 | 0.03 ± 0.00 |
| 12 | lSupramarginal Gyrus,<br>lSupramarginal Gyrus,<br>lSupramarginal Gyrus | (-48, -43, 26);<br>(-54, -46, 29);<br>(-54, -46, 35) | 0.00 ± 0.00 | 0.03 ± 0.01 |
| 13 | rPostCentral Gyrus,<br>rSupramarginal Gyrus | (51, -28, 56);<br>(66, -25, 32) | 0.00 ± 0.00 | 0.02 ± 0.01 |
| 14 | rPrecentral | (54, 5, 41) | 0.00 ± 0.00 | 0.02 ± 0.00 |
| 15 | rPostCentral | (45, -37, 59) | 0.00 ± 0.00 | 0.01 ± 0.00 |
| 16 | rSupramarginal Gyrus,<br>rPostCentral Gyrus | (57, -25, 50);<br>(63, -16, 41) | 0.00 ± 0.00 | 0.00 ± 0.00 |
| 17 | lPrecentral Gyrus | (-57, 5, 5) | 0.00 ± 0.00 | 0.00 ± 0.01 |

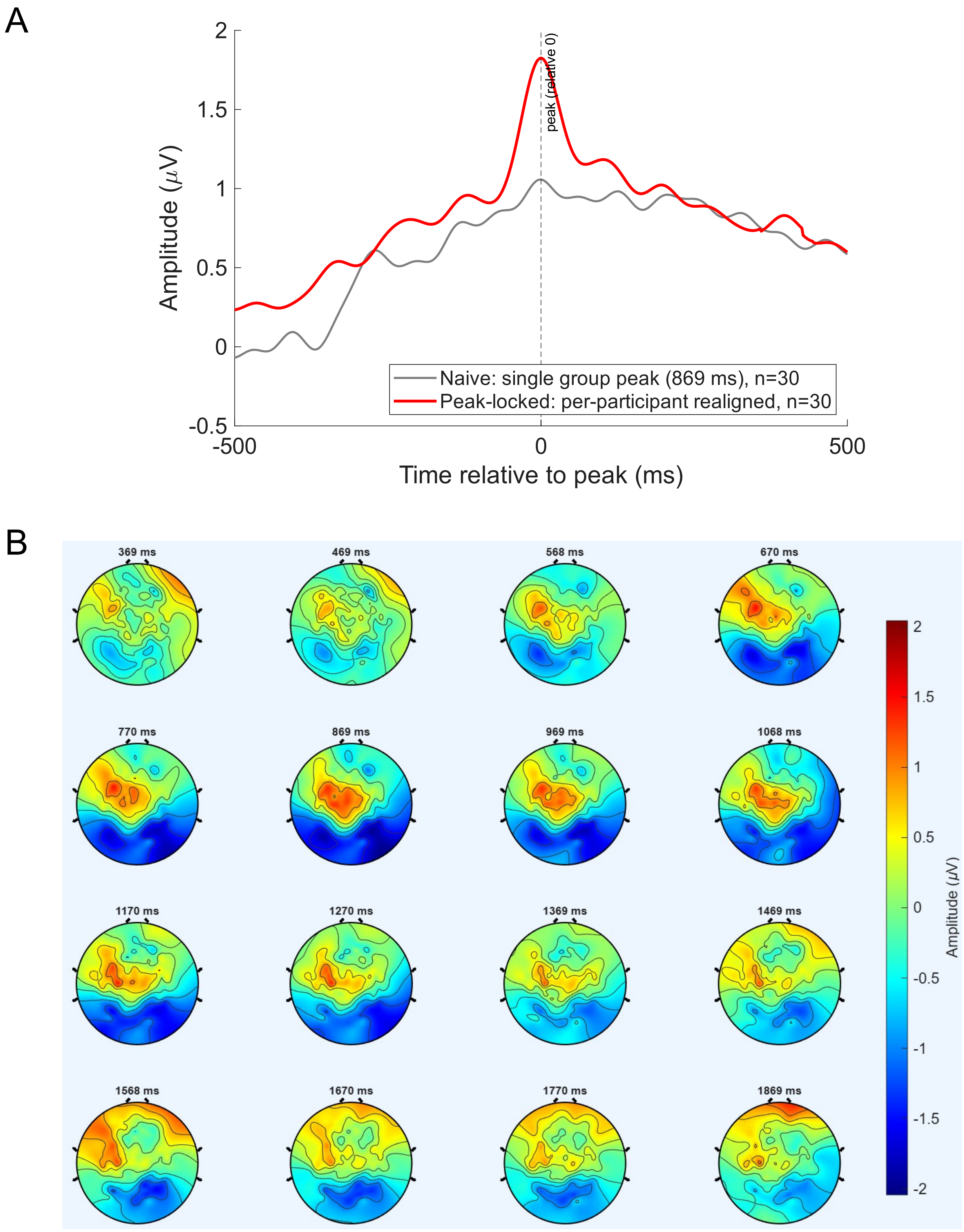


**Figure S14. The CPP during mental counting (no manual response).** (A) CPP-ROI waveform (A2, A3, A4, B2, D16) averaged after realigning each participant to their own peak (red), compared with the grand average aligned to the single group peak (grey). The realigned average is shown for visualisation only. The grand-average CPP, pooled over counting-up and counting-down blocks, peaked at 869 ms after target onset (counting down, 1127 ms; counting up, 859 ms). The forward-model fits used seven timepoints from 100 ms before to 50 ms after this peak (769–919 ms). (B) Scalp topography from 500 ms before to 1000 ms after the peak, in 100-ms steps, on a shared colour scale. All data were low-pass filtered at 10 Hz (zero-phase).

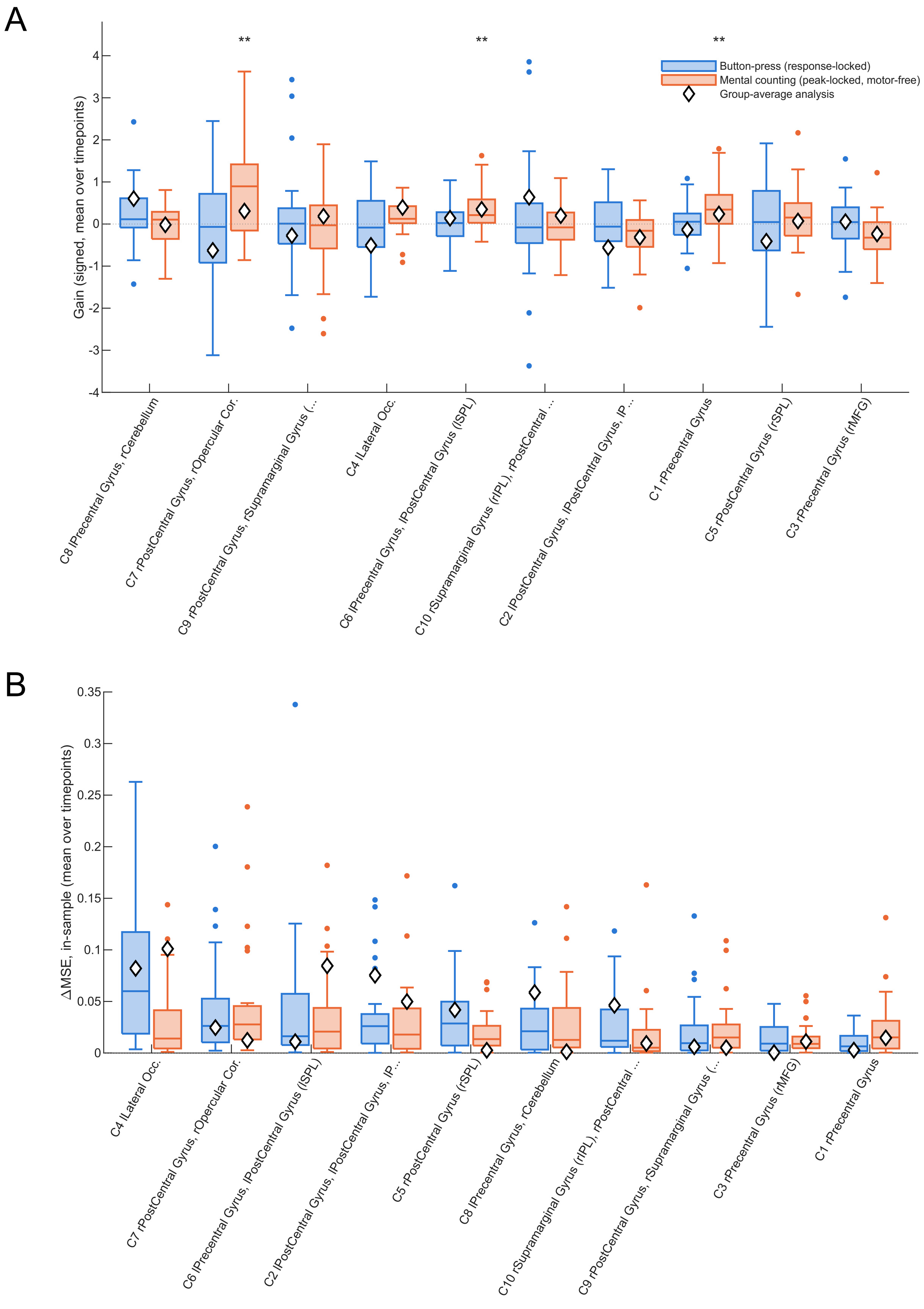

A
**
**
**
Button-press (response-locked)
Mental counting (peak-locked, motor-free)
Group-average analysis
Gain (signed, mean over timepoints)
C8 lPrecentral Gyrus, rCerebellum
C7 rPostCentral Gyrus, rOpercular Cor.
C9 rPostCentral Gyrus, rSupramarginal Gyrus (...
C4 lLateral Occ.
C6 lPrecentral Gyrus, lPostCentral Gyrus (lSPL)
C10 rSupramarginal Gyrus (rIPL), rPostCentral ...
C2 lPostCentral Gyrus, lPostCentral Gyrus, lP ...
C1 rPrecentral Gyrus
C5 rPostCentral Gyrus (rSPL)
C3 rPrecentral Gyrus (rMFG)
B
ΔMSE, in-sample (mean over timepoints)
C4 lLateral Occ.
C7 rPostCentral Gyrus, rOpercular Cor.
C6 lPrecentral Gyrus, lPostCentral Gyrus (lSPL)
C2 lPostCentral Gyrus, lPostCentral Gyrus, lP ...
C5 rPostCentral Gyrus (rSPL)
C8 lPrecentral Gyrus, rCerebellum
C10 rSupramarginal Gyrus (rIPL), rPostCentral ...
C9 rPostCentral Gyrus, rSupramarginal Gyrus (...
C3 rPrecentral Gyrus (rMFG)
C1 rPrecentral Gyrus

**Figure S15. Participant-level cluster contributions to the button-press and mental-counting CPP.** The forward model was fitted separately for each participant, using that participant's own FEM leadfield and scalp topographies. Button press (blue; response-locked, −100 to +50 ms; N = 31) and mental counting (orange; locked to the pooled CPP peak at 869 ms, no manual response; N = 29) were each analysed at seven timepoints. The model used 10 source clusters (Table S5), obtained at a correlation threshold of 0.69, the most permissive threshold at which every participant's design had a condition number below 30 (median 11, range 6.6–20; 24 raw sources: median 214). Mean in-sample normalised MSE was 0.38 for button press and 0.48 for mental counting.

(A) Signed cluster gain averaged over the seven timepoints, one value per participant, ordered by participant-level button-press gain rank. Asterisks mark clusters whose gain differed from zero across participants in a sign-flip permutation test with max-t family-wise error correction across clusters (10,000 permutations; **pFWER < .01). All three significant effects occurred in mental counting: C7 (right opercular/postcentral; pFWER = .004), C6 (pFWER = .003) and C1 (pFWER = .003). No button-press cluster reached significance. Ranking clusters by mean |gain|, with 95% rank intervals from 10,000 bootstrap resamples of participants, C7 ranked first in mental counting (interval 1–1) and second in button press (interval 1–9).

(B) Unique contribution of each cluster, measured as the increase in normalised MSE when that cluster is removed and the model refitted (in-sample leave-one-cluster-out ΔMSE). Values are averaged over timepoints, one per participant, and ordered by participant-level button-press ablation rank. C7 ranked first in mental counting (interval 1–4) and second in button press (interval 2–6), with a median ΔMSE of ≈0.03 in both conditions. Ablation rankings agreed between conditions (Spearman ρ = 0.75, bootstrap 95% CI 0.10–0.87), whereas gain rankings did not (ρ = −0.28, CI −0.71 to 0.50). In a paired permutation test on each cluster's share of total ΔMSE, no cluster contributed more with a manual response. Only C1 differed, contributing more during mental counting (uncorrected p = .045). When gains were estimated on N − 1 participants and ΔMSE was evaluated on the held-out participant (leave-one-subject-out), no cluster made a significant out-of-sample contribution (all pFWER > .05).

In both panels, boxes show the median and interquartile range, whiskers extend to 1.5 × IQR, dots mark outlying participants, and diamonds give the corresponding value from the fit to the group-average topography. Labels list the anatomical locations of member sources; C7 comprises the right opercular cortex (48, 2, 5) and right postcentral gyrus (60, −16, 23). All data were low-pass filtered at 10 Hz (zero-phase).

**Table S5. Composition of the 10 source clusters used in the participant-level analysis.** The 24 fMRI conjunction peaks were grouped by hierarchical clustering (average linkage on 1 − |r| between their forward-projected scalp topographies) at a correlation threshold of 0.69. Clusters therefore group sources by the similarity of their scalp projections, not by anatomical proximity. Short labels correspond to the axis labels in Fig. S15.

| Cluster | Short label | Member sources, MNI (x, y, z) |
|---|---|---|
| C1 | rPreCG | R precentral gyrus (63, 8, 8) |
| C2 | lPoCG/lSMG | L postcentral gyrus (−63, −19, 26; −63, −13, 23; −54, −13, 41; −45, −37, 32); L supramarginal gyrus/IPL (−63, −31, 23; −48, −43, 26; −54, −46, 29; −54, −46, 35) |
| C3 | rPreCG (MFG) | R precentral gyrus/MFG (54, 5, 41) |
| C4 | lLOC | L lateral occipital cortex (−54, −67, 5) |
| C5 | rPoCG (SPL) | R postcentral gyrus/SPL (45, −37, 59) |
| C6 | lPreCG/lSPL | L precentral gyrus (−51, −13, 53); L postcentral gyrus/SPL (−48, −40, 56) |
| C7 | rPoCG/rOperc | R postcentral gyrus (60, −16, 23); R opercular cortex (48, 2, 5) |
| C8 | lPreCG/rCereb | L precentral gyrus (−57, 5, 5); R cerebellum (24, −58, −19) |
| C9 | rPoCG/rSMG | R postcentral gyrus (51, −28, 56; 63, −16, 41); R supramarginal gyrus/IPL (57, −25, 50; 66, −25, 32) |
| C10 | rSMG/rPoCG | R supramarginal gyrus/IPL (63, −37, 26); R postcentral gyrus (48, −31, 59) |

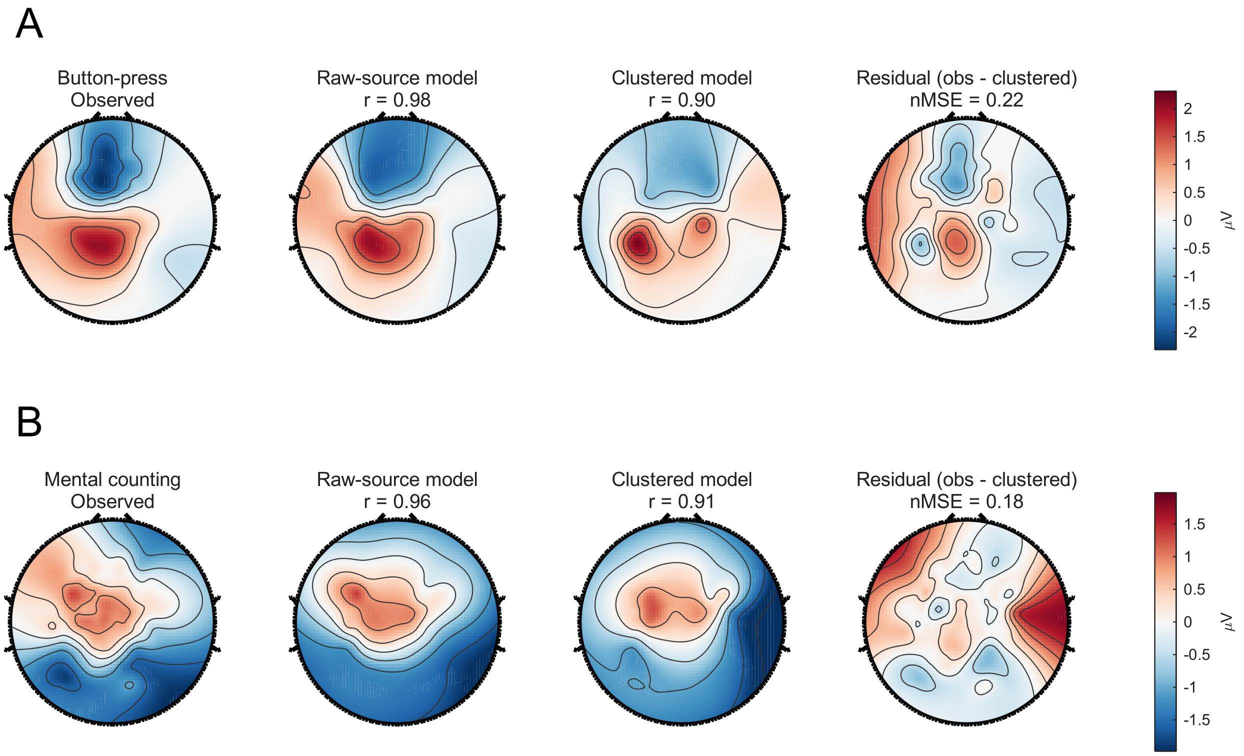


**Figure S16. Observed and model-predicted grand-mean topographies.** (A) Button press and (B) mental counting. From left to right, each row shows the observed grand-mean topography over the analysis window, the prediction of the 24-source (raw) model, the prediction of the 10-cluster model and the residual (observed − clustered model). Values above the model maps give the spatial correlation (r) with the observed topography; the value above the residual is the normalised MSE of the clustered fit. Each row shares one colour scale (μV). Both models used the participant-averaged, surface-normal-oriented, strength-free forward projections of the fMRI conjunction peaks, and the 10-cluster model used the threshold (0.69) required for well-conditioned participant-level fits. Across individual participants, the median spatial correlation between the observed topography and the participant's own clustered fit was r = 0.80 for button press and 0.73 for mental counting. Data were low-pass filtered at 10 Hz (zero-phase).
